# Therapeutic resistance in acute myeloid leukemia cells is mediated by a novel ATM/mTOR pathway regulating oxidative phosphorylation

**DOI:** 10.1101/2022.02.14.480468

**Authors:** Hae J. Park, Mark A. Gregory, Vadym Zaberezhnyy, Andrew Goodspeed, Craig T. Jordan, Jeffrey S. Kieft, James DeGregori

## Abstract

While leukemic cells are susceptible to various therapeutic insults, residence in the bone marrow microenvironment typically confers protection from a wide range of drugs. Thus, understanding the unique molecular changes elicited by the marrow is of critical importance towards improving therapeutic outcomes. In the present study, we demonstrate that aberrant activation of oxidative phosphorylation serves to induce therapeutic resistance in FLT3 mutant human AML cells challenged with FLT3 inhibitor drugs. Importantly, our findings show that AML cells are protected from apoptosis following FLT3 inhibition due to marrow-mediated activation of ATM, which in turn up-regulates oxidative phosphorylation via mTOR signaling. mTOR is required for the bone marrow stroma-dependent maintenance of protein translation, with selective polysome enrichment of oxidative phosphorylation transcripts, despite FLT3 inhibition. To investigate the therapeutic significance of this finding, we tested the mTOR inhibitor everolimus in combination with the FLT3 inhibitor quizartinib in primary human AML xenograft models. While marrow resident AML cells were highly resistant to quizartinib alone, the addition of everolimus induced profound reduction in tumor burden and prevented relapse. Taken together, these data provide a novel mechanistic understanding of marrow-based therapeutic resistance, and a promising strategy for improved treatment of FLT3 mutant AML patients.

## INTRODUCTION

Acute myeloid leukemia (AML) is an aggressive blood cancer with a high relapse rate and resistance to cytotoxic therapies (1). Internal tandem duplication (ITD) mutations in FMS-like tyrosine kinase 3 (FLT3) are among the most prevalent mutations in AML and are particularly associated with a poor prognosis (2). FLT3-ITD leads to constitutive activation of FLT3, a receptor tyrosine kinase, which induces activation of multiple effector molecules involved in survival, proliferation, and cell growth (3, 4). Clear scientific and clinical evidence that support the significance of activated FLT3 in leukemogenesis (5) has led to the development of several FLT3-targeted inhibitors.

Several clinical trials with two highly potent and selective second-generation FLT3 inhibitors, quizartinib and gilteritinib, have demonstrated their effectiveness in refractory AML patients with activating FLT3 mutations, showing significantly higher overall survival and rate of remissions compared to salvage chemotherapy (6–10). However, those remissions were short-lived. Furthermore, both FLT3 inhibitors showed noticeably delayed responses in the bone marrow (over weeks to a few months) compared to the peripheral blood (a few days), often showing persistence of the FLT3-ITD mutation in recovering marrow cells (11).

Indeed, it has long been appreciated that multiple components of the bone marrow (BM) microenvironment promote leukemogenesis, drug resistance, and relapse (12, 13). An extensive number of studies have shown that BM stromal cells (BMSCs) provide protection to leukemia cells from chemotherapies either through direct cell-to-cell contact or secretion of soluble factors (14). In particular, several recent studies have attempted to unravel the key human BMSC-activated signaling pathways in FLT3-ITD AML cells that lead to resistance to FLT3 inhibitors. In these studies, reactivation of STAT5 signaling and/or the MAPK/ERK pathway by human BMSCs were found to mediate the resistance to FLT3 inhibitors (15–18), yet conflicting data between each study indicates that the drug resistance mechanism is complex and likely to be cell-type dependent. Thus, further studies are needed to better understand the underlying mechanism of resistance to FLT3 inhibitors.

In this study, we uncover a novel pathway mediated by ATM and mTOR whereby BMSCs protect FLT3-ITD AML cells from apoptosis following FLT3 inhibition. We demonstrate that this ATM/mTOR pathway plays an essential role in the maintenance of protein translation and oxidative phosphorylation, which is critical for cell survival. Our data show that targeting these key mediators in collaboration with FLT3 inhibitor effectively overcomes BMSC-mediated protection and enhances elimination of AML cells.

## RESULTS

### Conditioned media of human bone marrow stromal cells prevents apoptosis in FLT3-ITD AML cells upon FLT3 inhibition

To study the paracrine effects mediated by BMSC, we utilized conditioned media from HS-5 human BM stromal cells (hBMSC-CM). The HS-5 cell line has been shown to be a reliable model to reproduce the biological properties mediated by BM mesenchymal stromal cells (19). Human AML cells either cultured with conditioned media from HS-5 or co-cultured with HS-5 exhibit enhanced resistance to multiple therapies including chemotherapy, immunotherapy, and targeted therapy (20–23). To further examine the role of hBMSC-CM on FLT3-ITD AML cells following FLT3 inhibition, we used MOLM-13 and MV4-11 cell lines. A significantly lower percentage of apoptotic cells were observed in both cell lines post-treatment with quizartinib in the presence of hBMSC-CM, compared to regular RPMI media (Fig. 1A). The protective effect of hBMSC-CM was also evident in the greater maintenance of viable cells observed after drug removal (Fig. 1B). In addition, we observed similar protection by hBMSC-CM for AML cells treated with gilteritinib (Supplementary Figs. S1A and S1B), another highly selective and potent FLT3 inhibitor.

**Figure 1.**
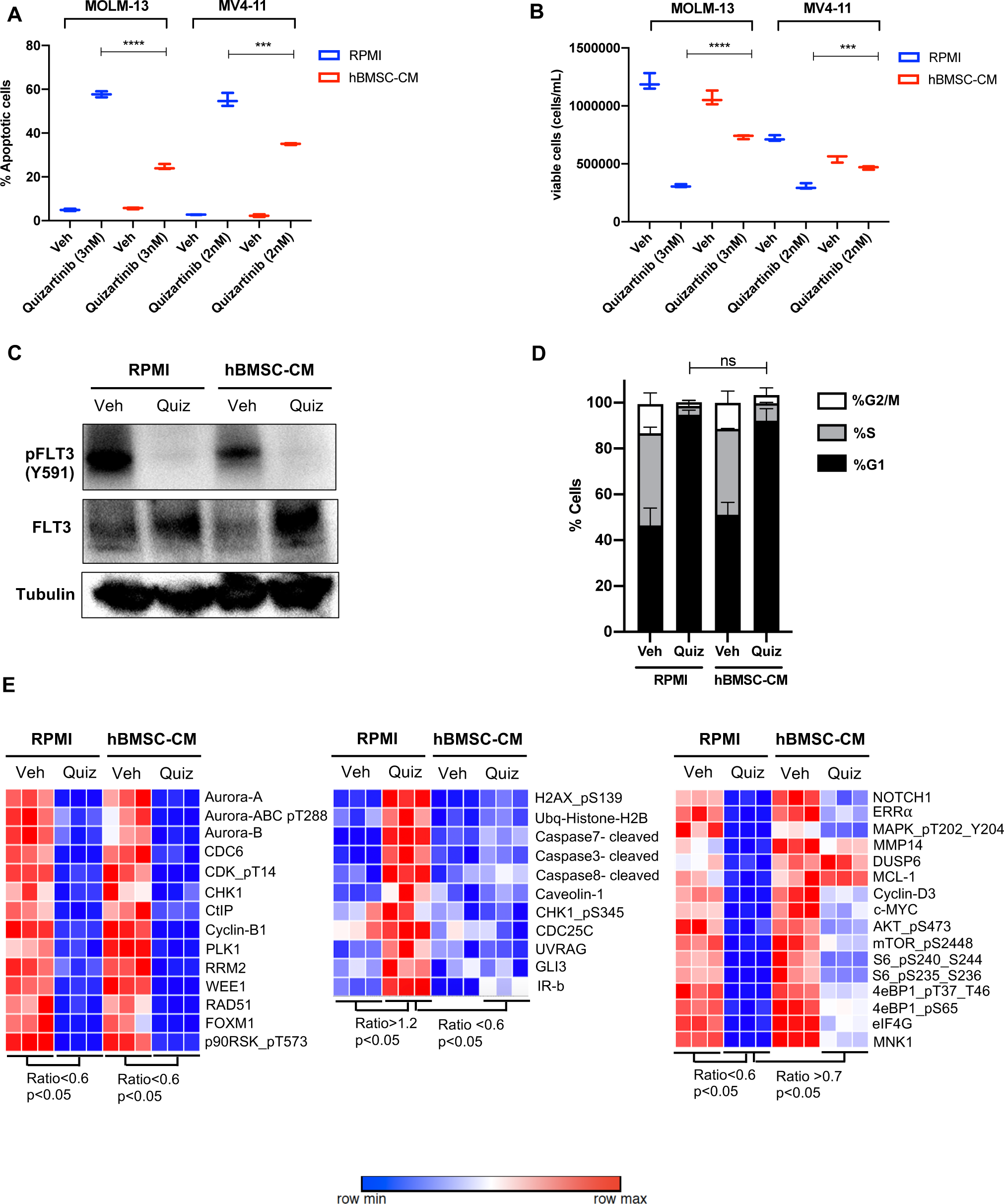
Conditioned media of human bone marrow stromal cells prevents apoptosis in FLT3-ITD AML cells upon FLT3 inhibition. **A** and **B**, FLT3-ITD AML cell lines, MOLM-13 and MV4-11, were treated with either DMSO (veh) or quizartinib with indicated doses for 48 hours in regular media (RPMI) or 50% conditioned media of human BM stromal cells in RPMI (hBMSC-CM). **(A)** Apoptotic cells were detected by 7-AAD/Annexin V staining post-treatment. **(B)** Cell viability was measured based on PI exclusion after 5 days of rebound growth post-treatment (n=3). **C**, MOLM-13 cells were harvested after 20 hours of indicated treatments (Quiz: 3nM quizartinib) to determine FLT3 activation by Western blotting. **D**, MOLM-13 cells were treated with the indicated conditions for 12 hours, followed by PI staining and cell cycle analysis via flow cytometry. **E**, MOLM-13 cells were harvested after 16 hours of treatments as indicated and subjected to reverse phase protein array analysis to explore the expression/activation levels of a wide range of proteins.

Next, we measured the activity of FLT3 in cells treated with quizartinib in either regular RPMI media or hBMSC-CM. Quizartinib completely inhibited the activity of FLT3 even in the presence of hBMSC-CM (Fig. 1C), suggesting that the drug resistance mediated by hBMSC- CM is not due to increased drug efflux or other mechanism preventing FLT3 inhibition. To examine if hBMSC-CM induces differential regulation of cell cycle upon FLT3 inhibition, we next measured cell cycle profiles in FLT3-ITD AML cells upon FLT3 inhibition in either RPMI or hBMSC-CM. Our cell cycle analysis demonstrated that cells surviving FLT3 inhibition show a similar G1 phase arrest, regardless of hBMSC-CM exposure (Fig. 1D).

Our results above indicate that the protective effect of hBMSC-CM is not due to prevention of FLT3 inhibition or alteration of cell cycle profiles. Hence, we further focused on hBMSC-CM-induced potential intrinsic changes that may promote cell survival following FLT3 inhibition. To explore alterations in protein expression and activation in FLT3-ITD AML cells caused by both FLT3 inhibition and hBMSC-CM exposure, we pursued a non-biased approach using reverse phase protein array (RPPA). We observed three major patterns of change: 1) downregulation upon FLT3 inhibition in both regular RPMI and hBMSC-CM (left panel, Fig. 1E), 2) upregulation with FLT3 inhibition in RPMI but not in hBMSC-CM (middle panel, Fig. 1E) or 3) downregulation in RPMI but not in hBMSC (right panel, Fig. 1E). We found that major components of the first pattern largely consist of cell cycle regulatory proteins (AURORA kinases, CDC6, CDK, CHK1, Cyclin B1, PLK1, and WEE1), consistent with our cell cycle analysis (Fig.1D). Regarding the second pattern of change (upregulation by FLT3 inhibition in RPMI but not in hBMSC-CM), proteins involved with the DNA damage response (p-H2AX, ubiquitinated histone H2b, p-CHK1, CDC25C, and UVRAG) or apoptosis (cleaved caspases 3, 7, and 8) were found to be in this category, consistent with our observations of differential induction of apoptosis. Finally, proteins involved in a wide range of biological functions were found to show a pattern of downregulation with FLT3 inhibition in RPMI but significantly less downregulation or nearly complete maintenance in hBMSC-CM. This group included the oncogenic transcription factor c-MYC, antiapoptotic protein MCL-1, and MAPK/ERK signaling proteins (DUSP6 and p-MAPK/ERK). Among these, MAPK/ERK signaling (16) and MCL-1 (24) are consistent with findings from other studies. Notably, multiple proteins involved in the mTOR signaling pathway (p-AKT, p-mTOR, p-S6, p-4E-BP1, eIF4G, and MNK1) demonstrate this consistent pattern. Together, the above data suggest that hBMSC-CM dramatically alters expression and activation of multiple pathways, and these changes may contribute to the survival of FLT3-ITD AML cells following FLT3 inhibition.

### Bone marrow microenvironment reverses mTOR and MYC pathway suppression in FLT3-ITD AML cells upon FLT3 inhibition *in vitro* and *in vivo*

To characterize the influence of the hBMSC-CM on gene expression in FLT3-ITD AML cells upon FLT3 inhibition, we compared the transcriptomes of MOLM-13 cells treated with either vehicle or quizartinib in RPMI or hBMSC-CM by performing RNA sequencing (RNA-seq). As shown in Fig. 2A, FLT3 inhibited AML cells in hBMSC-CM displayed a distinct gene expression signature compared to those in RPMI. Gene set enrichment analyses (GSEA) indicated that multiple pathways related to cell fate determination are significantly more enriched in AML cells in hBMSC-CM compared with RPMI after FLT3 inhibition (Fig. 2B). Specifically, a majority of mTOR complex1 (mTORC1) signaling genes and MYC target genes demonstrated significant suppression post-FLT3 inhibition in RPMI, while such downregulation was at least partially reversed in the presence of hBMSC-CM (Fig. 2C), further supporting our findings from the RPPA data. We observed that hBMSC-CM induced higher baseline mTOR activity and substantially prevented downregulation of mTOR activity by FLT3 inhibition (Fig. 2D and Supplementary table). Baseline c-MYC expression was also higher in hBMSC-CM compared to RPMI, and while complete suppression of c-MYC expression was observed at early timepoints after treatment with quizartinib in both conditions, c-MYC protein expression rebounded at later time points only in the presence of hBMSC-CM (Fig. 2D and Supplementary table).

**Figure 2.**
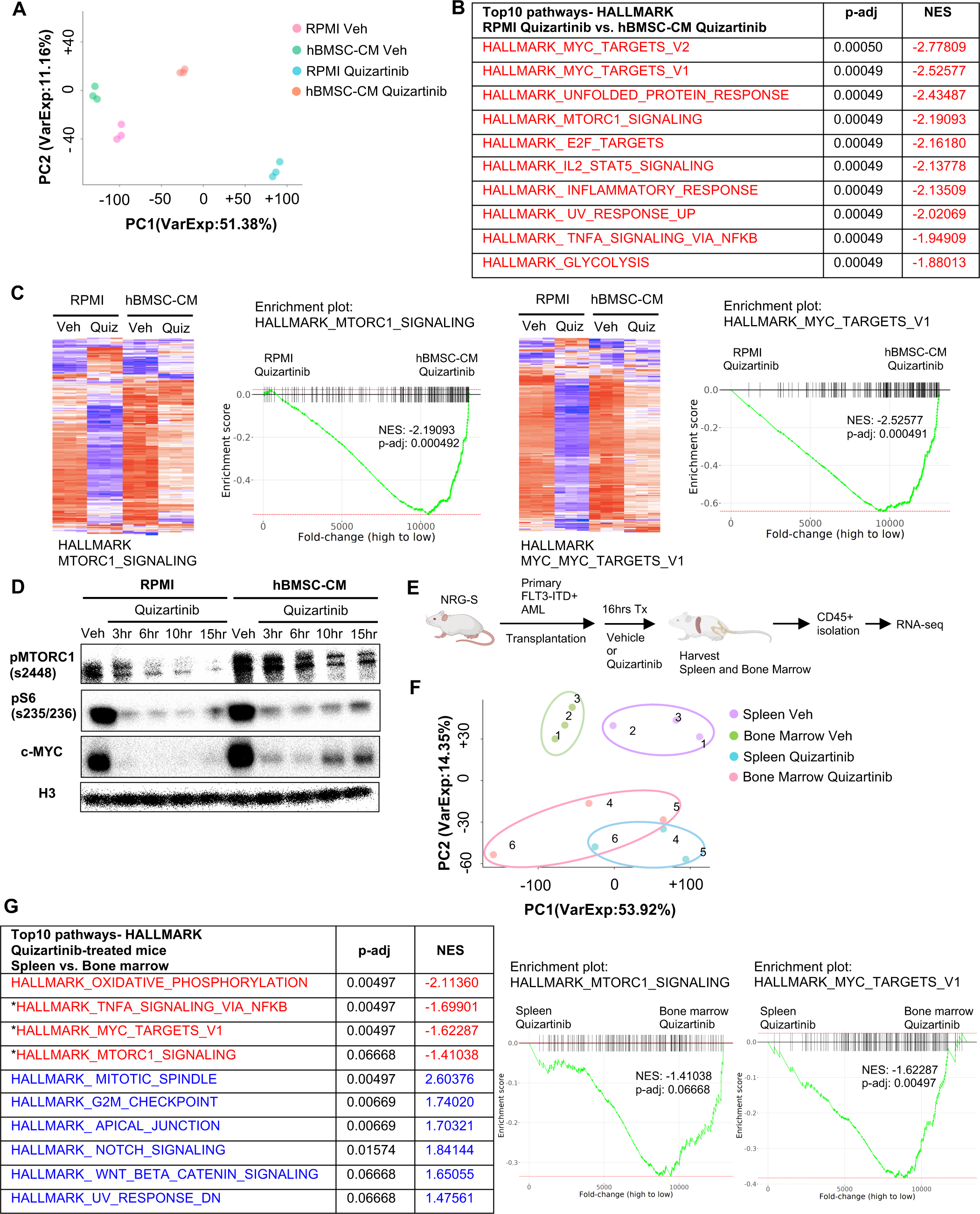
The bone marrow microenvironment reverses mTOR and MYC pathway suppression in FLT3-ITD AML cells upon FLT3 inhibition *in vitro* and *in vivo*. A-C, RNA from MOLM-13 cells was harvested after 15 hours of treatment with vehicle or 3nM quizartinib in either RPMI or hBMSC-CM and subjected to RNA-seq (n=3). (A) Principal Component Analysis (PCA) plot of each group. (B) Top 10 enriched signaling pathways (by p-adj) from GSEA of Hallmark gene sets applied on RNA-seq data from quizartinib-treated cells in hBMSC-CM compared with quizartinib-treated cells in RPMI. Pathways with significant alteration (p-adj < 0.1) and negative normalized enrichment score (NES) are represented with red. (C) Enrichment plots and heat maps showing comparisons of transcriptomes of mTORC1 signaling or MYC targets between quizartinib (Quiz) treated cells in hBMSC-CM and quizartinib-treated cells in RPMI. D, MOLM-13 cells were treated as indicated and harvested at different timepoints (3, 6, 10, and 15 hours) to measure mTORC1 signaling and c-MYC protein levels by Western blotting. E, Design of *in vivo* study using human AML xenograft mouse model. NRG-S mice were transplanted with primary FLT3-ITD^+^ AML cells. One cohort of mice was treated with vehicle, while the other cohort was treated with 2.5 mg/kg quizartinib for 16 hours (n=3). After spleen and BM were harvested, human AML cells were isolated by CD45 isolation kit and RNA from each sample were collected for RNA-seq. F, PCA plot of RNA-seq data from human AML samples isolated from spleen and BM. Samples from the same mouse were denoted by the same number. G, Top 10 enriched signaling pathways (by p-adj) from GSEA of Hallmark gene sets applied on RNA-seq data from human AML cells. Samples from the BM of quizartinib-treated mice were compared with the samples from the spleen of the same paired mouse, and top 10 pathways are shown on the left. Pathways with significant alteration (p-adj < 0.1) are represented with colors (Red: negative NES, Blue: positive NES). Asterisks denote pathways that exhibited similarly significant enrichment patterns from *in vitro* study (Fig. 2B). Enrichment plots of mTORC1 signaling or MYC target genes are shown on the right.

To explore the *in vivo* relevance of our findings, we performed a transcriptomic analysis of patient-derived primary FLT3-ITD^+^ AML cells engrafted in NOD.*Rag1^-/-^;γc^null^* (NRG) mice expressing human cytokines GM-CSF, IL-3, and SCF (NRG-S mice). We chose to use NRG-S mice for our human xenograft mouse model because they not only express human cytokines previously shown to provide partial protection from FLT3 inhibition, but also demonstrate efficient engraftment of several patient-derived AML cells (25). To investigate the effect of the BM microenvironment on gene expression profiles of human AML cells, we compared transcriptomes of AML cells isolated from the spleen and the BM of mice treated with vehicle or quizartinib for 16 hours (Fig. 2E). Cells isolated from the spleen showed a distinct gene expression signature compared to those from the BM, and treatment of quizartinib resulted in significant alterations of transcriptome profiles both in the spleen and the BM (Fig. 2F). Our GSEA data showed multiple pathways with a similar gene expression pattern between AML cells from the spleen of quizartinib-treated mice and AML cells treated with quizartinib in RPMI, indicating similar effects of quizartinib on gene expression in both our *in vitro* and *in vivo* models (Supplementary Fig. S2A). Next, we examined if the BM microenvironment of NRG-S mice recapitulates the transcriptomic changes we observed in hBMSC-CM from our *in vitro* studies. Indeed, our GSEA data revealed that expression of mTORC1 signaling genes and MYC target genes were significantly more enriched in AML cells isolated from the BM than those from the spleen of quizartinib-treated mice, consistent with findings from our *in vitro* model (Figs. 2C and 2G). Notably, genes of these pathways did not show significant differences between the BM and spleen of vehicle-treated mice, suggesting higher expression of mTORC1 signaling and MYC target genes in the BM is specific to the context of FLT3 inhibition (Supplementary Fig. S2B). In contrast, other pathways such as Oxidative Phosphorylation and G2/M Checkpoint exhibited a similar enrichment in the BM compared to spleen for both vehicle and quizartinib treated mice (Fig. 2G and Supplementary Fig. S2C).

Collectively, our *in vitro* and *in vivo* models demonstrate consistent gene expression patterns, indicating that the BM microenvironment promotes maintenance of mTOR signaling and MYC target gene signatures in FLT3-ITD AML cells following FLT3 inhibition.

### Targeting the mTOR pathway reverses bone marrow mediated protection of FLT3-ITD AML cells from FLT3 inhibition

As a downstream effector of activated FLT3 kinase, mTOR signaling has been implicated in the survival of FLT3-ITD^+^ AML cells, and aberrant activation of the mTOR pathway has been reported in FLT3-ITD AML cell lines that developed intrinsic resistance to FLT3 inhibitors (26, 27). Our findings from unbiased studies using RPPA (Fig. 1E) and RNA-seq (Fig. 2) also suggest the potential role of the mTOR pathway in the BM-mediated survival of FLT3-ITD AML cells upon FLT3 inhibition. We first determined whether pharmacological inhibition of mTOR signaling reverses hBMSC-CM-mediated protection from apoptosis following FLT3 inhibition.

Our data revealed that while the mTOR inhibitor everolimus alone did not significantly promote apoptosis, combinatorial treatment of everolimus with quizartinib significantly reversed hBMSC-CM-mediated protection from apoptosis mediated by quizartinib (Fig. 3A).

**Figure 3.**
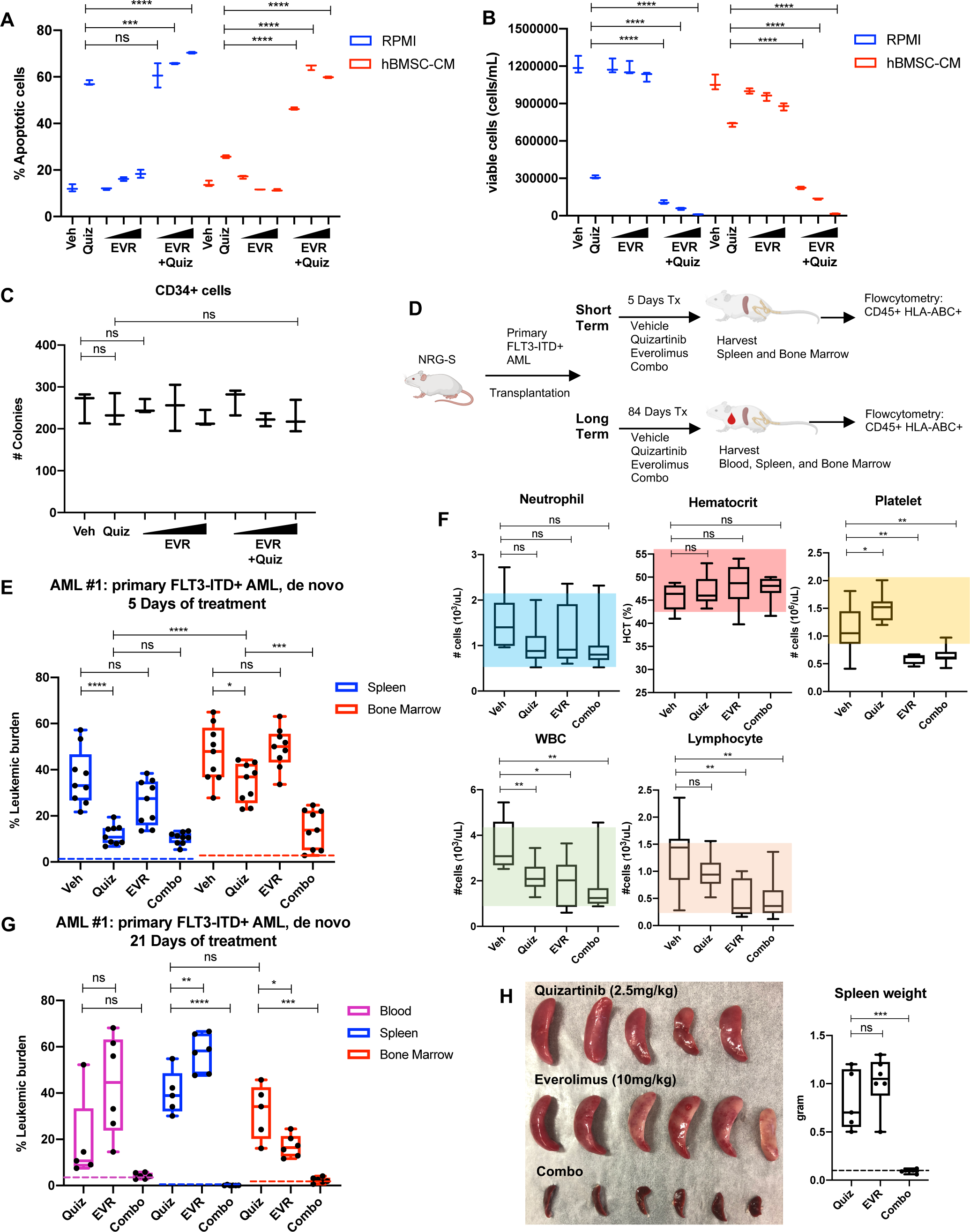
Targeting the mTOR pathway reverses the bone marrow mediated protection of FLT3-ITD AML cells from FLT3 inhibition. **A** and **B**, MOLM-13 cells were treated with 3nM quizartinib (Quiz) and everolimus (EVR) alone or together as indicated (increasing concentrations of everolimus of 5, 10, and 50nM indicated by triangles) for 48 hours in RPMI or hBMSC-CM (n=3). (**A**) Apoptosis was measured post-treatment, and (**B**) cell viability was measured after 5 days of rebound growth post-treatment. **C**, CD34^+^ cord blood cells from a healthy newborn were treated with 10nM quizartinib and everolimus alone or together as indicated (increasing concentrations of everolimus of 10, 50, and 100nM indicated by triangles) for 48 hours. 25,000 cells from each group were seeded in methylcellulose post-drug treatments. After 11 days, the number of colonies were counted (n=3). **D**, Schematic diagram of experimental design. NRG-S mice were transplanted with human primary FLT3-ITD^+^ AML (AML #1: a 69-year-old female patient who were newly diagnosed AML with FLT3-ITD and NPM1 mutations). When leukemic burden reached 2-5% (∼21-28 days after transplantation, determined by measuring CD45^+^ HLA-ABC^+^ cells in the blood), mice were divided into two groups. For short-term treatment, mice were treated with 1) vehicle, 2) 2.5mg/kg quizartinib, 3) 10mg/kg everolimus, 4) combo for 5 days, daily p.o. (n=9). For long-term treatment, mice were treated with the same regimen for 84 days, daily p.o. (n=7-10). After treatment, leukemic burden was measured by quantifying HLA-ABC^+^ CD45^+^ cells via flow cytometry in spleen, BM, and blood (for long-term treatment only). **E**, Plot showing leukemic burden of short-term treatment group; dotted line represents negative controls from non-transplanted mice. **F**, Complete Blood Count (CBC) from blood samples of mice from long-term treatment group at day 5 of treatment were plotted. Colored boxes represent reference values (neutrophil: 0.54- 2.15x10^3^ cells/uL, hematocrit: 44.1-57.2%, platelet:0.914-2.055 x10^3^ cells/uL, WBC: 0.94- 4.12x10^3^ cells/uL, lymphocyte: 0.26-1.56x10^3^ cells/uL). **G**, Plot showing leukemic burden of long-term treatment group, dotted line represents negative controls from non-transplanted mice. **H**, Photograph of spleens from long-term treatment group on the day of harvest. Spleen weight from each group (dotted line represents average spleen weight from healthy untreated mice) is shown plotted on the right.

Furthermore, AML cells treated with quizartinib/everolimus in hBMSC-CM failed to recover after removal of drugs, in contrast to cells treated with quizartinib alone (Fig. 3B). Similar results were observed in FLT3-ITD MV4-11 cells (Supplementary Figs. S3A and S3B).

To assess whether normal human hematopoietic progenitors would be affected by the combination therapy, we treated CD34^+^ selected cord blood cells from a healthy newborn with quizartinib and/or everolimus, followed by plating on methylcellulose for clonogenic assays.

None of these treatments showed significant changes in colony numbers, suggesting minimal effects on healthy hematopoietic progenitors (Fig. 3C).

To test whether inhibition of mTOR could enhance the efficacy of quizartinib in eliminating FLT3-ITD^+^ AML cells *in vivo*, we tested the combination therapy in our xenograft mouse model. As illustrated in Fig. 3D, mice engrafted with patient-derived FLT3-ITD^+^ AML cells were treated with vehicle, quizartinib or everolimus alone, or these drugs in combination for 5 days to assess the short-term effects of these therapies on leukemic burden in the spleen and BM. A separate cohort of mice were treated with the same drug regimen for 84 days to examine the long-term effects of the therapies. After 5 days of treatment, quizartinib alone substantially reduced AML cells in the spleen, but only modestly reduced AML burden in the BM (Fig. 3E).

On the other hand, the addition of everolimus to quizartinib (“combo” therapy) resulted in effective elimination of leukemia burden in the BM, recapitulating our *in vitro* data with hBMSC- CM. Complete blood counts revealed that mice treated with everolimus alone or in combination with quizartinib showed a modest level of thrombocytopenia and leukopenia without affecting neutrophils or hematocrit (Fig. 3F), consistent with patient data that everolimus has been reported to induce hematologic changes that are tolerable (28). Mice on the long-term treatments did not exhibit reductions in weight (Supplementary Fig. S3C). While mice receiving long-term treatment with quizartinib alone eventually relapsed with high leukemic burden detected in the blood, BM and spleen, leukemia cells were not detected in any compartment from mice treated with the combination therapy (Fig. 3G). Elimination of the leukemia in the spleen by the combination therapy was further confirmed by the absence of splenomegaly (Fig. 3H). Furthermore, in an independent experiment with a second primary FLT3-ITD+ AML sample from a patient who developed relapse after chemotherapy, we observed potent elimination of leukemic burden in all compartments with the combination therapy. While monotherapy with quizartinib was effective in peripheral tissues (spleen and blood), effective elimination of leukemia cells in the bone marrow was only achieved with the combinatorial treatment of quizartinib with everolimus (Supplementary Fig. S3D). As before, we did not observe significant alterations in mouse weights, indicating that the therapy was well tolerated (Supplementary Fig. S3E).

In all, these data show that mTOR pathway plays a key role in the BM-mediated protection of FLT3-ITD AML cells from FLT3-targeted therapy.

### Restoration of mTOR signaling by hBMSC-CM limits downregulation of protein translation upon FLT3 inhibition

Control of protein translation is known to be one of the critical proliferation pathways mediated by mTOR signaling in cancer cells (29). In particular, regulation of protein synthesis via mTOR signaling has been implicated in human myeloid leukemogenesis, making it an attractive potential therapeutic target in AML cells (30). Therefore, we speculated that altered protein translation would be an important downstream effect of mTOR signaling that drives the BM-mediated survival of FLT3-ITD^+^ AML cells following FLT3 inhibition. Indeed, our transcriptome analysis revealed significantly higher expression of genes involved in translation regulation in cells treated with quizartinib in hBMSC-CM than those in RPMI (Fig. 4A). Moreover, FLT3 inhibition in RPMI dramatically reduced phosphorylation of 4E-BP1 and ribosomal protein S6 that regulate translation downstream of mTORC1 signaling, while addition of hBMSC-CM led to partial maintenance of these activities (Fig. 4B), consistent with the partial restoration of mTOR activity. Importantly, dual inhibition of mTOR and FLT3 eliminates the effect of hBMSC- CM on maintenance of these translation regulators.

**Figure 4.**
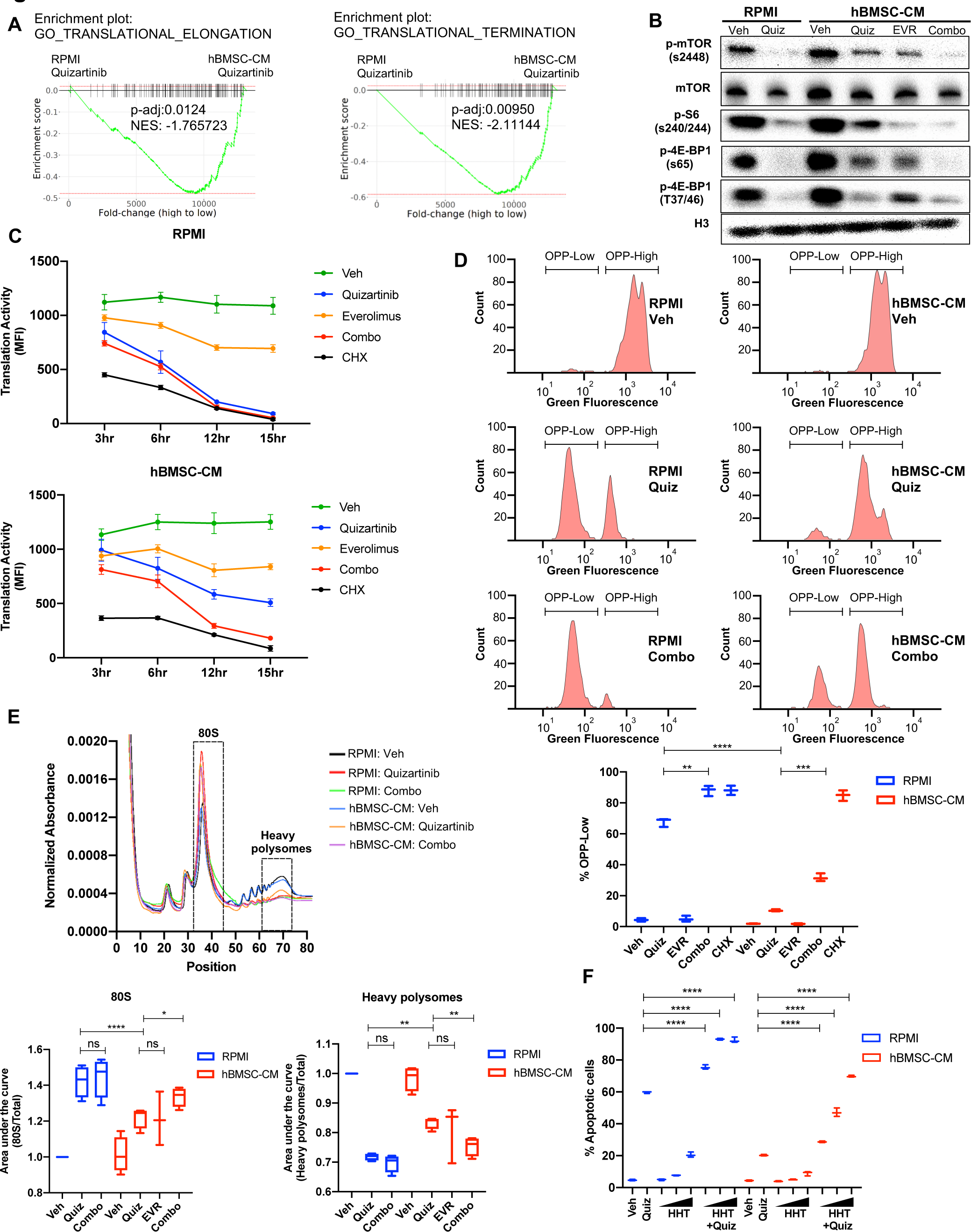
Restoration of mTOR signaling by hBMSC-CM limits downregulation of protein translation upon FLT3 inhibition. **A**, GSEA of Gene Ontology gene sets from *in vitro* study (Fig. 2 A-C) showing enrichment of translational control in AML cells treated with quizartinib in hBMSC-CM, compared with quizartinib-treated cells in RPMI. **B**, MOLM-13 cells were treated with vehicle, 3nM quizartinib, 10nM everolimus, or quizartinib+everolimus (combo) in RPMI or hBMSC-CM for 15 hours and harvested for Western blotting to measure mTOR signaling and its downstream mediators involved in translational control. **C**, MOLM-13 cells were treated with the indicated drugs at different timepoints (3, 6, 12, and 15 hours), followed by labeling with o-propargyl-puromycin (OPP), to measure translation activity using flow cytometry (MFI: mean fluorescence intensity for OPP incorporation). Cells were treated with 10ug/mL cycloheximide (CHX) as a control for inhibition of translation (n=3). **D**, Representative flow cytometry histograms of OPP assay from cells treated with vehicle, quizartinib, and combo in RPMI or hBMSC-CM for 15 hours, showing binary peaks (OPP-High and OPP-Low). Percent of OPP- Low population from all treatment groups is shown at the bottom. **E**, MOLM-13 cells were treated with the indicated drugs for 12 hours and harvested for polysome profiling. Area under curve for each treatment was normalized to the vehicle-treated group in RPMI and are shown for 80S peak fraction (position 33-44) and heavy polysome fraction (position 62-73) at the bottom (n=3 for everolimus treated cells in hBMSC-CM, n=4 for all other groups). **F**, MOLM-13 cells were treated with 3nM quizartinib and homoharringtonine (HHT) alone or together as indicated (increasing concentrations of HHT of 1, 5, and 10nM indicated by triangles) for 48 hours in RPMI or hBMSC-CM, followed by measurement of apoptosis (n=3).

To directly assess effects on protein translation, we measured translation activity in FLT3-ITD AML cells after each drug treatment by detecting cells labeled with O-propargyl-puromycin (OPP), a puromycin analogue that is incorporated into newly translated proteins, allowing a reliable quantification of protein translation (31). Strikingly, FLT3 inhibition alone was sufficient to dramatically reduce translation activity in FLT3-ITD AML cells over the course of treatment, comparable to the effect of translation inhibition by cycloheximide, but the reduction was significantly limited by hBMSC-CM. Moreover, combined inhibition of mTOR and FLT3 reversed the partial restoration of translation activity in hBMSC-CM, indicating that the effect of hBMSC-CM on translation activity is dependent on restored mTOR activity (Fig. 4C). The number of viable cells was maintained in every treatment condition through this timecourse, indicating that the significant reduction in translation activity was not due to concurrent cell death (Supplementary Fig. S4A). Following FLT3 inhibition, a substantial population of AML cells showed a very low level of OPP labeling (OPP^low^) in RPMI, indicating severe suppression of translation, while hBMSC-CM significantly reduced the proportion of OPP^low^ cells and the OPP^high^ cells showed a noticeable leftward peak shift compared to vehicle-treated cells (consistent with a partial inhibition of translation on a per cell basis). Furthermore, dual inhibition of mTOR and FLT3 by combination treatment resulted in dramatic increase of OPP^low^ cells even in the presence of hBMSC-CM (Fig. 4D). Our data from earlier timepoints suggest that leftward peak shift in OPP^high^ cells and accumulation of OPP^low^ cells occur over the course of drug treatment (Supplementary Fig. S4B). Notably, mTOR inhibition alone showed a leftward shift in OPP^high^ cells without noticeable detection of OPP^low^ cells (Supplementary Fig. S4C), consistent with a partial reduction in overall translation (that apparently is well-tolerated by cells).

To determine how hBMSC-CM affects translation of specific mRNAs in FLT3-ITD AML cells following inhibition of FLT3 and/or mTOR signaling, we performed polysome fractionization using ultracentrifugation through sucrose density gradients followed by analysis of mRNA distribution profiles. As expected, cells treated with quizartinib in RPMI showed a much higher proportion of transcripts bound to the 80S monosome and substantially reduced polysome-associated transcripts, consistent with impaired translation activity (Fig. 4E). In contrast, FLT3 inhibition in hBMSC-CM showed partial restoration of polysome content, especially heavy polysomes that consist of 5 or more ribosomes, with a concomitant decrease in the 80S monosome peak. Moreover, addition of everolimus to quizartinib treatment reversed the partial restoration of heavy polysomes in hBMSC-CM. Findings from our polysome profiling are consistent with what we observed from OPP assays, further validating the role of restored mTOR signaling by hBMSC-CM in regulation of protein translation. Finally, we tested if pharmacological inhibition of translation recapitulates the effect of mTOR inhibition in reversing hBMSC-CM-mediated protection from apoptosis following FLT3 inhibition. Indeed, combinatorial treatment of quizartinib with the translation inhibitor homoharringtonine (HHT) showed a significantly higher number of apoptotic cells compared to cells treated with quizartinib alone, even in the presence of hBMSC-CM (Fig. 4F), phenocopying the effect of everolimus. These results further support previously reported synergistic elimination of FLT3- ITD AML cells following treatment with FLT3 inhibitors in combination with HHT (32). Taken together, our data suggest that mTOR-dependent partial restoration of translation activity is a key survival mechanism of FLT3-ITD AML cells in hBMSC-CM following FLT3 inhibition.

### Examination of polysome enrichment of transcripts reveals a critical role for oxidative phosphorylation in hBMSC-CM-mediated protection from FLT3 inhibition

Our OPP assays and polysome profiling data indicate hBMSC-CM induces partial restoration of translation activity and polysome-associated transcripts following FLT3 inhibition, respectively. Yet, the mechanism for how cells survive despite incomplete restoration of translation remains unknown. To investigate this, we further analyzed mRNAs enriched in heavy polysomes to examine if specific subsets of mRNAs were selectively translated. First, we measured levels of RNAs isolated from the whole lysates (input) and heavy polysome fractions. As expected, cells treated with quizartinib alone or quizartinib and everolimus in combination demonstrated a significant reduction of RNA levels that was substantially more extensive in polysomes than inputs (Fig. 5A), consistent with the suppressive effects of FLT3 inhibition on polysome engagement of transcripts. mRNAs isolated from input and heavy polysomes were subjected to RNA-seq. Notably, GSEA applied on RNA-seq data from input (unfractionated) samples indicated that the addition of everolimus to quizartinib in hBMSC-CM significantly suppressed a majority of the pathways that were shown to be restored by hBMSC- CM following treatment of quizartinib alone, including genes for mTORC1 signaling, MYC targets, oxidative phosphorylation, and translation regulation (Figs. 5B and 5C; compare to Figs. 2B and 2G). Thus, mTOR signaling is essential for the ability of hBMSC-CM to maintain the expression of these key pathways in FLT3-inhibited cells.

**Figure 5.**
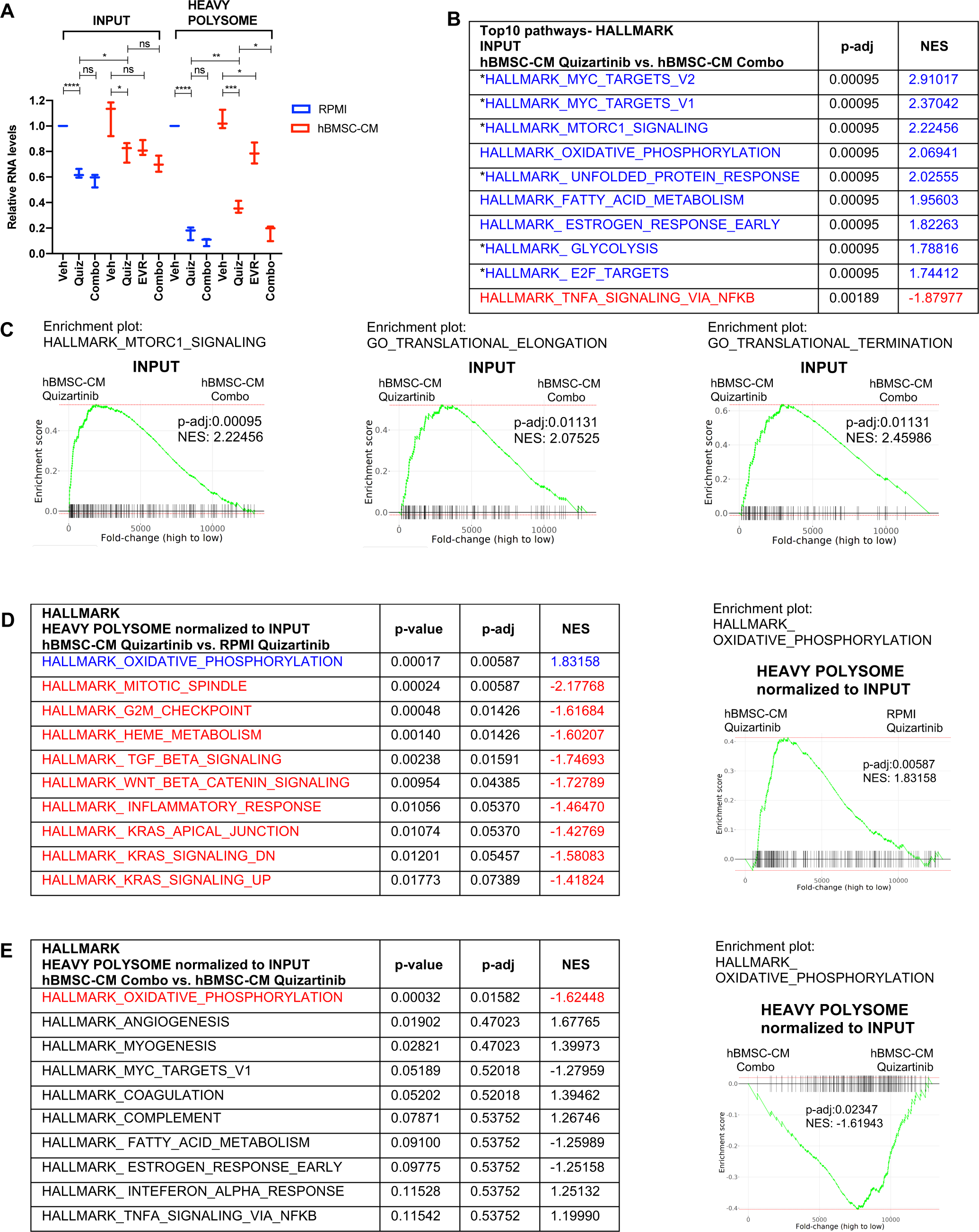
Polysome enrichment of transcripts by hBMSC-CM following FLT3 inhibition is highly selective for oxidative phosphorylation mRNAs and is mTOR-dependent. A, RNA yields from whole cell lysates (input) and heavy polysome fractions Fig. 4E) were measured and normalized to vehicle-treated group in RPMI for each biological replicate. B and C, GSEA of Hallmark and Gene Ontology gene sets was performed on RNA-seq data from input samples. Quizartinib-treated cells in hBMSC-CM were compared with combo-treated cells in hBMSC-CM. (B), Top 10 enriched signaling pathways from GSEA of Hallmark gene sets with significant alterations (p-adj < 0.1) represented with colors (Red: negative NES, Blue: positive NES). Asterisks denote pathways which exhibited partial restoration following quizartinib treatment in hBMSC-CM, compared to those in RPMI (Fig. 2B). (C), Enrichment plots of mTOR Signaling, Translational Elongation, and Translational Termination are shown. D and E, Top 10 enriched signaling pathways (by p-adj) from GSEA of Hallmark gene sets for RNA-seq data from human AML cells from heavy polysome associated mRNAs normalized to input. Pathways with significant alteration (p-adj < 0.1) are represented with colors (Red: negative NES, Blue: positive NES). Analyses were done by comparing (D) cells treated with quizartinib in hBMSC-CM with those in RPMI or (E) cells treated with combo in hBMSC-CM with those treated with quizartinib alone in hBMSC-CM. Enrichment plots of Oxidative Phosphorylation pathway for each comparison are shown on the right.

In order to identify transcripts that are preferentially retained or depleted from polysomes, we performed GSEA on mRNAs in heavy polysomes normalized to input. Remarkably, only one Hallmark pathway, oxidative phosphorylation, exhibited significant enrichment in heavy polysomes isolated from cells treated with quizartinib in hBMSC-CM, compared to those in RPMI (Fig. 5D). Furthermore, combinatorial treatment of quizartinib with everolimus significantly reduced polysome enrichment of oxidative phosphorylation mRNAs, even in the presence of hBMSC-CM, compared to quizartinib alone (Fig. 5E). In fact, oxidative phosphorylation was the only Hallmark gene set with significant (by p-adj) alteration with the combination treatment. Interestingly, as shown in Fig. 2G, oxidative phosphorylation was also the most enriched pathway identified in primary AML cells in the BM relative to the spleen in mice treated with quizartinib. Our GSEA of Hallmark gene sets demonstrate that other pathways with statistical significance showed opposite enrichment patterns, suggesting polysome enrichment of transcripts by hBMSC-CM following FLT3 inhibition is highly selective to oxidative phosphorylation mRNAs and is mTOR-dependent (Figs. 5D and 5E). In accordance with this, GSEA of Gene Ontology gene sets demonstrated similar polysome enrichment patterns for transcripts that are involved with mitochondria integrity and biogenesis (Supplementary Fig. S5).

To examine whether altered expression and polysome enrichment of oxidative phosphorylation mRNAs translates into corresponding changes in mitochondrial respiration, we measured the oxygen consumption rate (OCR) of FLT3-ITD AML cells following inhibition of FLT3 and/or mTOR in either RPMI or hBMSC-CM. In agreement with our findings from GSEA, we found a dramatic decrease in OCR in cells following FLT3 inhibition, while the OCR reduction was partially restored in the presence of hBMSC-CM (Fig. 6A). Furthermore, the dual inhibition of FLT3 and mTOR reversed this partial restoration of OCR. Of note, the degree of restoration was greater for maximal respiration than basal respiration, as represented by reserve respiratory capacity which reflects the difference between maximal and basal respiratory rate (Fig. 6A). The observed changes in OCR correlate with similar alterations in ATP levels (Fig. 6B), indicating that restoration of OCR is able to rescue ATP, which may be key for cell survival. Indeed, combinatorial treatment of quizartinib with ATP synthase inhibitor oligomycin A significantly reversed hBMSC-CM-mediated protection from apoptosis (Fig.6C), consistent with the effect of everolimus or HHT (Figs. 3A and 4F), as well as our previous study that showed that tyrosine kinase inhibition engenders acute sensitivity to oligomycin A in myeloid leukemias (33).

**Figure 6.**
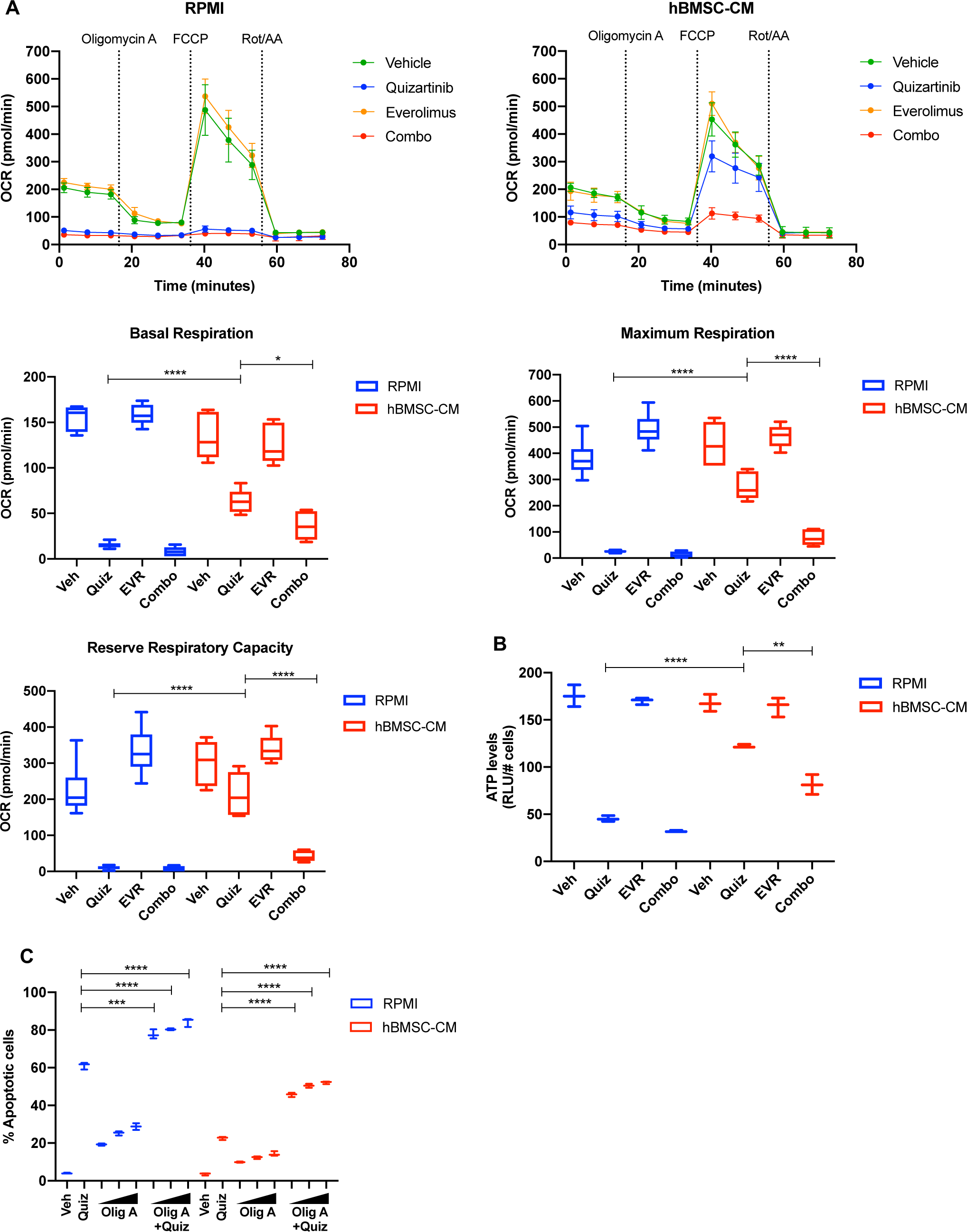
mTOR-dependent oxidative phosphorylation is a key survival mechanism of hBMSC-CM-mediated protection from FLT3 inhibition. **A**, MOLM-13 cells were treated with indicated drugs for 15 hours, followed by measurement of oxygen consumption rate (OCR) by Seahorse XF Cell Mito Stress Test (n=5). Basal respiration, maximum respiration, and reserve respiratory capacity are shown at the bottom. **B**, MOLM-13 cells were treated with indicated drugs for 15 hours, followed by luminescence-based ATP assays (normalizing Relative Light Units [RLU] to cell number). **C**, MOLM-13 cells were treated with 3nM quizartinib and Oligomycin A (Olig A) alone or together as indicated (increasing concentrations of Olig A of 1, 2, and 4 nM indicated by triangles) for 48 hours in RPMI or hBMSC-CM, followed by measurement of apoptosis (n=3).

Taken together, our data suggest that the polysome enrichment of oxidative phosphorylation mRNAs mediates survival of FLT3-ITD AML cells in hBMSC-CM following FLT3 inhibition through an mTOR-dependent mechanism, potentially by maintenance of energy levels.

### ATM mediates the protection by hBMSC-CM upstream of mTOR signaling and oxidative phosphorylation

Having established as axis between mTOR activity and oxidative phosphorylation, we next sought to better understand factors upstream of mTOR that may regulate activity critical to survival of AML cells. Our previous work has shown that inhibition of ATM (ataxia telangiectasia mutated) in FLT3-ITD AML cells causes synergistic cell killing with FLT3 inhibitors by inducing apoptosis through exacerbation of mitochondrial oxidative stress (34). Additionally, there has been accumulating evidence that ATM drives the BM-mediated resistance to chemotherapy in other hematologic malignancies, such as multiple myeloma and acute lymphoblastic leukemia via upregulation of cytokines such as IL-6 (35) or CCL3, CCL4, and GDF15 (36). We found that blocking FLT3 inhibited ATM expression in RPMI, but that ATM expression was maintained in hBMSC-CM following FLT3 inhibition in FLT3-ITD AML cells (Fig. 7A). However, dual inhibition of mTOR and FLT3 did not reverse the effect of hBMSC-CM on ATM levels, suggesting ATM acts upstream of mTOR. We observed that phosphorylated ATM (S1981) was predominantly detected at a smaller size (∼250KDa) than full-length ATM (350KDa) following FLT3 inhibition in RPMI, while hBMSC-CM significantly restored full-length phosphorylated ATM (Fig.7A and Supplementary table). Moreover, dual inhibition of mTOR and FLT3 resulted in a higher ratio of small to full-length phospho-ATM. Treatment of cells with pan-caspase inhibitor Z-VAD-FMK resulted in the elimination of the small ATM isoform following FLT3 inhibition, suggesting that cleavage of full-length ATM was mediated by caspase activity (Supplementary Fig. S6A), as previously reported from other studies (37, 38). Co-treatment of Z-VAD-FMK with quizartinib in RPMI partially rescued cell death (Supplementary Fig. S6B).

**Figure 7.**
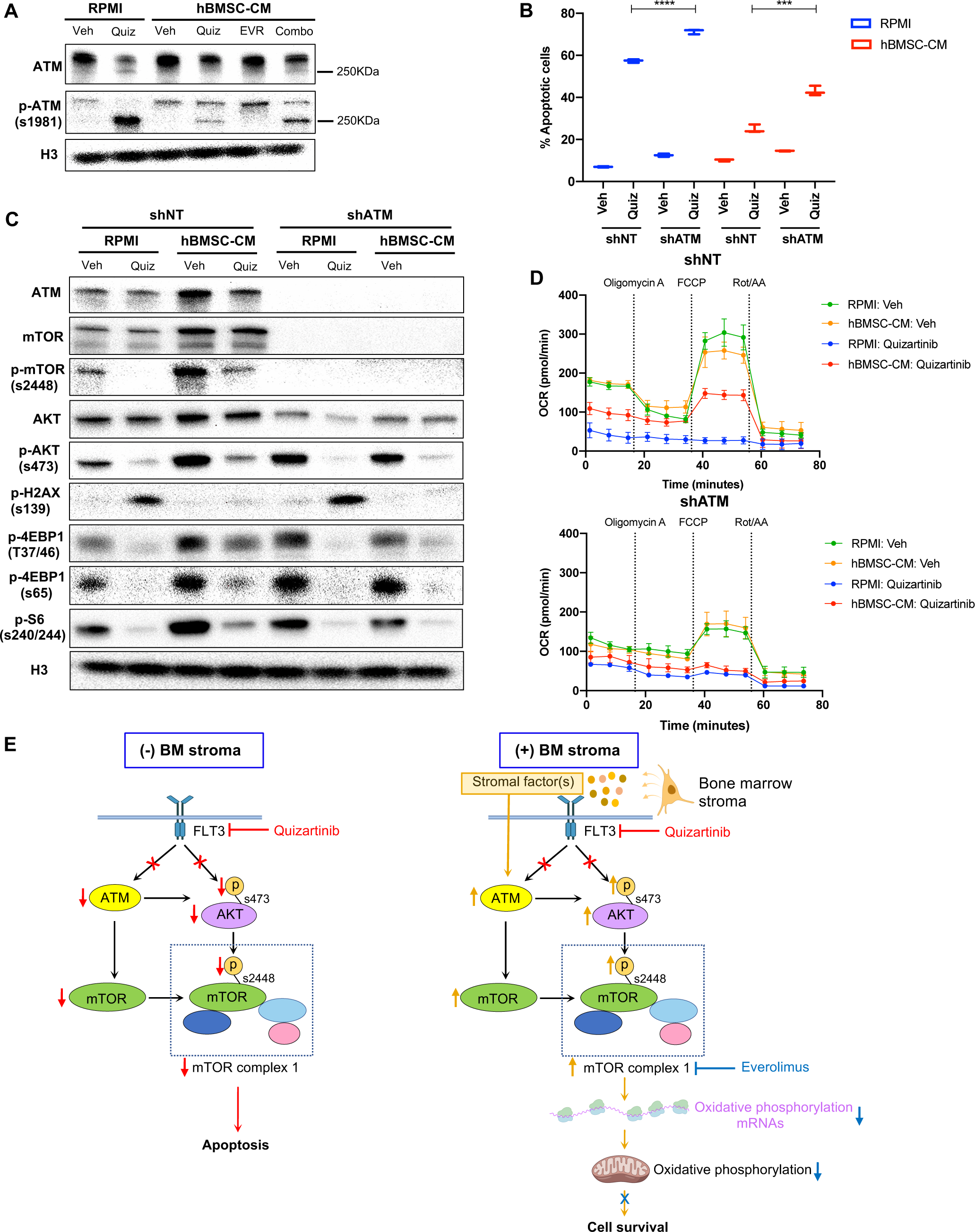
ATM mediates the protection from FLT3 inhibition by hBMSC-CM upstream of mTOR signaling and oxidative phosphorylation. **A**, MOLM-13 cells were treated with vehicle, 3nM quizartinib, 10nM everolimus, and combo in RPMI or hBMSC-CM for 15 hours and harvested to determine levels of total and phosphorylated ATM by Western blotting. **B**, MOLM-13 cells were transduced with lentiviruses expressing shRNAs (shNT: non-targeting shRNA or shATM: ATM-targeting shRNA). Cells were treated with either vehicle or 3nM quizartinib (Quiz) for 48 hours in RPMI or hBMSC-CM, followed by measurement of apoptosis (n=3). **C**, MOLM-13 cells expressing shNT or shATM were treated with the indicated conditions for 15 hours and harvested for Western blot analyses. **D**, MOLM-13 cells expressing shNT or shATM were treated with the indicated conditions for 15 hours, followed by measurement of OCR by Seahorse XF Cell Mito Stress Test (n=4). **E**, Model of ATM and mTOR-dependent survival of FLT3-ITD AML cells following FLT3 inhibition in BM microenvironment. In the absence of BM stroma, FLT3-ITD AML cells undergo apoptosis upon FLT3 inhibition as a result of downregulation of ATM and mTOR complex 1 (mTORC1) activity (left). In the presence of BM stroma, on the other hand, FLT3-ITD AML cells are protected from cell killing effects by FLT3 inhibition (right). As indicated by orange arrows, the BM stroma-mediated protection is driven by ATM through regulation of mTOR protein levels, as well as mTORC1 activity via AKT, subsequently leading to partial restoration of translation. Restored mTORC1 activity selectively drives translation of oxidative phosphorylation mRNAs that promote cell survival even in the presence of FLT3 inhibition. The BM stroma-mediated protection against apoptosis upon FLT3 inhibition is reversed with combinatorial treatment with everolimus, which inhibits mTOR signaling and downstream processes including polysome enrichment and selective translation of oxidative phosphorylation mRNAs, and oxidative phosphorylation. Effects of drugs that target FLT3 (quizartinib) or mTOR (everolimus) are represented with red or blue, respectively, arrows and X’s.

Next, we examined the potential role of ATM in hBMSC-CM-mediated protection of FLT3-ITD AML cells following FLT3 inhibition. Compared to cells expressing non-targeting shRNA (shNT), cells with shRNA-mediated ATM knockdown (shATM) demonstrated reversal of protection against apoptosis upon FLT3 inhibition in hBMSC-CM (Fig. 7B), as well as impaired recovery of cell growth after removal of quizartinib (Supplementary Fig. S6C). Given that ATM has long been known to regulate cell cycle checkpoints, we asked if knocking down ATM allowed cell cycle progression despite FLT3 inhibition via bypassing ATM-dependent checkpoint arrest. We found that shATM did not prevent the G1 arrest observed with quizartinib with or without hBMSC-CM (Supplementary Fig. S6D), indicating that the mechanism of hBMSC-CM-mediated protection by ATM is not through activating cell cycle checkpoints.

Based on the importance of mTOR signaling in hBMSC-CM-mediated protection, we wanted to further investigate the potential association between ATM and mTOR signaling in FLT3-ITD AML cells. Strikingly, knockdown of ATM resulted in the complete absence of detectable mTOR protein expression (validated with two different antibodies as described in Methods), without significantly affecting the baseline phosphorylation of 4E-BP1 and S6 (Fig. 7C). Still, upon FLT3 inhibition, shATM prevented the restoration of p-4E-BP1 or p-S6 otherwise elicited by hBMSC-CM. Additionally, shATM not only resulted in a moderate reduction of AKT protein levels, but also impaired the hBMSC-CM-mediated restoration of phosphorylation of AKT following FLT3 inhibition (Fig. 7C and Supplementary table). These results implicate ATM as a key regulator of mTORC1 signaling through regulation of the mTOR protein itself, as well as AKT that has been shown to regulate mTORC1 via phosphorylation (29). It is important to note that we observed the same results in CRISPR/Cas9-mediated ATM knockout FLT3-ITD AML cells (Supplementary Fig. S6E). The ATM knockout cells also showed greater sensitivity to FLT3 inhibition in RPMI and reversal of protection against apoptosis upon FLT3 inhibition in hBMSC-CM, compared to control cells (Fig. S6F), further validating our findings. In addition, we confirmed that in other AML cell lines, MV4-11 (FLT3-ITD), THP-1 or OCI-AML3 (both FLT3 WT), the degree of ATM knockdown correlated with the extent of reductions in mTOR expression and with AKT phosphorylation in MV4-11 and THP-1 (Supplementary Fig. S7A and Supplementary table). Furthermore, our analysis of data sets from Gene Expression Omnibus (GEO) indicates that brain tissues from human ataxia-telangiectasia (A-T) patients, who have mutated ATM, have significantly lower mTOR mRNA levels compared to control groups, coinciding with a reduction of gene expression associated with mTORC1 signaling and PI3K- AKT-mTOR pathways according to GSEA (Supplementary Fig. S7B). These data implicate that the role of ATM in controlling mTOR signaling extends to other cell types.

We next measured OCR in FLT3-ITD AML cells expressing shRNAs (shNT or shATM) following inhibition of FLT3 with or without hBMSC-CM. ATM knockdown resulted in a significant reduction of baseline OCR (Fig. 7D and Supplementary Fig. S7C). More importantly, ATM knockdown in combination with quizartinib prevented the partial restoration of OCR elicited by hBMSC-CM, consistent with our findings with dual inhibition of FLT3 and mTOR. Taken all together, our data indicate that ATM drives hBMSC-CM-mediated protection of AML cells from apoptosis following FLT3 inhibition through regulation of mTOR signaling and oxidative phosphorylation.

## DISCUSSION

Results presented here support a key role for a novel pathway mediated by ATM, mTOR, translational control, and oxidative phosphorylation, whereby BM stromal cells protect FLT3- ITD AML cells from therapeutic elimination following FLT3 inhibition (Fig. 7E). Multiple components of the hBMSC-CM-dependent pathway are essential for AML survival, as substantiated by the loss of protection when each mediator is targeted by pharmacological inhibition or genetic knockdown. Notably, this survival pathway can be effectively blocked at multiple points using FDA-approved drugs, including everolimus and HHT, which should facilitate the translation of these studies into clinical trials for FLT3-ITD AML. In all, these studies reveal the therapeutic potential of combining FLT3 inhibition with therapeutic targeting of mediators of this pathway to improve clinical outcomes for patients with FLT3-ITD AML.

Although numerous clinical trials have investigated the efficacy of FLT3 inhibitors both as monotherapy and in combination with other therapeutics for patients with FLT3-mutated AML, these trials have thus far not yielded prolonged remissions (11). This clinical challenge indicates that a better understanding of resistance mechanisms is necessary to identify additional targets for effective molecularly targeted AML therapy. The BM microenvironment has been shown to mediate resistance to FLT3 inhibitors via several mechanisms including: 1) high levels of FLT3 ligand in the BM microenvironment that leads to persistent activation of FLT3 (39); 2) cytochrome p450 enzyme produced by BMSCs enhances metabolization of FLT3 inhibitors (40); and 3) BMSCs-mediated activation of prosurvival pathways such as MAPK/ERK and/or STAT5 signaling (15–18). Additionally, a recent study has shown that the BM microenvironment is responsible for early resistance of FLT3-ITD+ AML cells to FLT3 inhibition, followed by clonal expansion of pre-existing NRAS mutant subclones that drive relapse (41). Collectively, these studies suggest that BM-mediated resistance against FLT3- targeted therapy in FLT3-dependent AML cells is multifactorial and context-dependent.

In our present studies, we focused on intrinsic changes in AML cells upon FLT3 inhibition in the presence of BM microenvironmental factors by utilizing FLT3-ITD AML cell lines and xenograft mouse models utilizing primary FLT3-ITD+ AML cells. Both our *in vitro* and *in vivo* data consistently support the conclusion that persistent activation of mTOR signaling mediated by the BM microenvironment impedes cell death induced by FLT3 inhibition. Notably, activation of mTOR signaling has been previously shown in FLT3-WT AML cells co-cultured with mouse BM stroma (42), suggesting upregulation of the mTOR pathway in AML cells may not be limited to dependence on FLT3 or human BM stromal factors. Moreover, our therapeutic studies in mice demonstrated the limitation of FLT3-targeted monotherapy in eliminating leukemia from the BM compared to the periphery (spleen and blood), leading to relapse even with sustained treatment. Our results indicate that such relapse can be effectively prevented by combination therapy that targets FLT3 and mTOR signaling. Given that everolimus in combination with midostaurin (a multi-kinase inhibitor that targets FLT3) was well tolerated in patients with AML in a phase 1 clinical study (43), our results suggest a promising treatment strategy for patients with FLT3-dependent AML.

Over the past two decades, many aspects of mTOR’s role in controlling key cellular processes, including protein synthesis, have been extensively elucidated, and mechanistic details of mTOR-mediated pathways in different cellular and physiological contexts remain an area of active study (44). Whereas mTOR signaling has long been known to control global protein synthesis, studies have shown that it can also drive the selective translation of specific subsets of mRNAs through mechanisms that remain elusive (45–48). Our data reveal a novel process through which mTOR-dependent translational control of transcripts involved in oxidative phosphorylation is a mechanism by which BMSCs mediate survival of FLT3-ITD AML cells following FLT3 inhibition (Fig. 7E). The mechanism we propose consolidates some of the major findings from studies that explore the role of mTOR signaling and oxidative phosphorylation in different physiological contexts: 1) Enhanced mTORC1 signaling is required for translation of mitochondria-related mRNAs during erythropoiesis (47); 2) mTORC1 controls mitochondrial respiration and biogenesis via regulation of 4E-BPs in human breast cancer cells (48); 3) Upregulation of oxidative phosphorylation confers chemoresistance via inhibition of AMPK pathway and subsequent activation of mTORC1 signaling in FLT3-WT AML cells co-cultured with hBMSC (49).

Recent studies have shown that drug-resistant AML cells and leukemia stem cells are highly dependent on oxidative phosphorylation for energy production, leading to clinical trials of several drugs that target mitochondrial metabolism, thus directly/indirectly inhibiting oxidative phosphorylation (50, 51). Compared to normal hematopoietic cells or solid tumors, AML cells have been shown to have low reserve respiratory capacity, rendering them highly sensitive to inhibition of oxidative phosphorylation and oxidative stress (52). Our data suggest that the significant restoration of maximal respiration rate and reserve respiratory capacity by hBMSC- CM may promote cell survival by preventing ATP depletion following FLT3 inhibition. While combination therapy effectively reversed the hBMSC-CM-mediated restoration of oxidative phosphorylation and ATP, mTOR inhibition alone did not significantly affect oxidative phosphorylation and ATP levels, suggesting that FLT3-ITD signaling may activate alternative pathway(s) to maintain mitochondrial bioenergetics. Given that low reserve respiratory capacity is a specific metabolic vulnerability of AML cells, we speculate that therapeutic targeting of the mTOR/translation/oxidative phosphorylation axis could block BM-mediated resistance to other AML therapies.

One of the novel aspects of our findings is the unconventional role of ATM in regulation of mTOR signaling and thus oxidative phosphorylation. While many regulators of the mTOR pathway have been identified in a wide range of cellular contexts, the control of the kinase activity of the two major mTOR complexes, mTORC1 and mTORC2, remains a primary focus to understand regulatory inputs of mTOR signaling (29, 44). Yet, the regulation of mTOR protein levels is poorly understood. We discovered that ATM is required for the expression of the mTOR protein, independent of FLT3 activity or BM stromal effects. It is important to note that ATM-mediated regulation of mTOR protein levels does not appear to be unique to FLT3- dependent AMLs or leukemia in general, as implicated with our analyses of other cell types. Furthermore, our data suggest that ATM not only regulates mTOR protein levels, but also mediates the phosphorylation/activity of AKT. As ATM knockdown only reduced AKT activity in the context of FLT3 inhibition, ATM and FLT3 appear to redundantly regulate AKT activity.

Given that AKT has been identified as a downstream effector of FLT3 signaling (3, 4), BM stromal factors likely confer dependence of FLT3-ITD AML cells on ATM following FLT3 inhibition to maintain AKT activity, followed by subsequent activation of mTORC1. Similarly, our observation that S6 and 4E-BP1 were still phosphorylated under baseline condition even in the absence of mTOR in ATM deficient cells suggests that other determinants of S6 and 4E- BP1 activity beyond mTOR exist, perhaps involving other FLT3-dependent signaling molecules.

Based on these intriguing findings, it will be of interest to elucidate the mechanism of ATM-mediated regulation of mTOR signaling. Indeed, ATM has been shown to mediate numerous cellular homeostasis pathways, independent of its functions in response to DNA damage. ATM has been shown to be involved in insulin signaling by promoting protein translation by phosphorylating 4E-BP1 in HEK293 cells and adipose tissue (53), and by activating AKT via its kinase function to enhance glucose uptake in muscle cells (54). In contrast to what we show here, ATM has been shown to *inhibit* mTOR signaling in response to oxidative stress (55) or hypoxia (56), implicating diverse and context-dependent functions of ATM.

In summary, our findings demonstrate that the BM microenvironment provides a unique survival pathway mediated by ATM, mTOR, and oxidative phosphorylation in FLT3-ITD AML cells following FLT3 inhibition. Therapeutic targeting of this pathway, such as by inhibition of mTOR, in combination with FLT3 inhibition could lead to more durable remissions for patients with FLT3-ITD+ AML.

## MATERIALS AND METHODS

### Cell Lines and Cell Culture

All human AML cells (MOLM-13, MV4-11, THP-1, OCI-AML3) were cultured in RPMI medium supplemented with 10% FBS and 1% antibacterial antimycotic (anti-anti). Other non-AML cells (SUP-B15) were cultured in IMDM medium supplemented with 10% FBS and 1% anti-anti. The adherent cell lines 293FT was grown in DMEM with 10% FBS and 1% anti-anti. All the cell lines were authenticated by short tandem repeat examination and tested negative for Mycoplasma using the e-Myco plus PCR Detection Kit (iNtRON) in October 2021. All transfections were performed in 293FT cells using FuGENE HD Transfection Reagent (#E2311; Promega) at 3:1:1:1 ratio of shRNA (shNT: #SHC016 or shATM: #TRCN0000039948 in pLKO.1-puro; MilliporeSigma) or CRISPR-Cas9:sgRNA constructs (sgNT:GAGCTGGACGGCGACGTAAA or sgATM: TCTACCCCAACAGCGACATG in lentiCRISPR V2) and packaging plasmids pMDLg/pRRE, pMD.G, pRSV-Rev in OPTI-MEM solution (#31985070; Gibco). Viral supernatant was collected 36 and 48 hours after transfection. Spin infections were performed at room temperature at 1,500 × g for 1 hour with polybrene reagent (1:2,000; Fisher Scientific). For the generation of hBMSC-CM, HS-5 cells were cultured in RPMI medium supplemented with 10% FBS and 1% anti-anti. At 80-90% confluency, conditioned media was collected and filtered through 0.22μm PES sterile filter. hBMSC-CM was prepared by combining 50% of conditioned media of HS-5 with 50% RPMI medium supplemented with 10% FBS and 1% anti-anti.

### Isolation of CD34^+^ cells

Peripheral blood mononuclear cells (PBMC) were isolated from fresh human cord blood cells by density centrifugation on Ficoll-Paque, from which CD34+ cells were isolated by the MACS CD34 MicroBead kit (#130-046-702; Miltenyi Biotec). CD34^+^ cells were incubated in serum free media (IMDM supplemented with 20% BIT 9500 Serum Substitute, 10ng/mL SCF, 10ng/mL IL-3, and 10ng/mL FLT-3) overnight prior to drug treatments.

### Apoptosis and Cell viability assays

Cells were seeded at 1.0 to 2.0 x10^5^ /mL in triplicate wells of 48-well tissue culture plates. Where indicated, the cells were treated with drug(s) for 48 hours. After treatment, cells were stained using 7-aminoactinomycin D (7-AAD)/anti-Annexin V (Nexin reagent, MilliporeSigma) to detect apoptotic cells. For cell viability assay, a sample of cells from each well was reseeded at 40,000 /mL in RPMI media and incubated 5 days to measure rebound growth post-treatment. Cells were stained with propidium iodide (PI;10mg/mL) and viable cells (PI^-^) were counted with a flow cytometer (Millipore Guava easyCyte 8HT). Adenosine triphosphate (ATP) levels were measured using the Cell Titer-Glo Assay (Promega). ATP data were normalized to cell number.

### Translation activity measurements

*OPP Assay.* Cells were treated with the indicated conditions and labelled with 20µM OPP for 30 minutes. Protein synthesis was measured using Click-iT™ Plus OPP Alexa Fluor™ 488 Protein Synthesis Assay Kit (#C10456; Life Technologies) per manufacturer’s instructions.

Translation activity was quantified by measuring green fluorescence intensity with a flow cytometer (Millipore Guava easyCyte 8HT).

### Polysome Profiling

Prior to harvest, cells were treated with the indicated conditions, followed by treatment with 100μg/mL cycloheximide for 10 minutes to halt translation. 3x10^7^ cells were harvested from each treatment group. Following harvesting, cells were resuspended in lysis buffer (20mM HEPES pH 7.4, 15mM MgCl2, 200mM NaCl, 1% Triton x- 100, 0.1mg/mL cycloheximide, 2mM DTT, 100U/mL RNasin ribonuclease inhibitor). Cell homogenates were spun down for 5 min at 13,000 x g to pellet any cellular debris, then 500 µL of this clarified lysate was loaded on 10-60% sucrose gradients in SW41 tubes in lysis buffer lacking Triton X-100. These gradients were prepared using a BioComp system and chilled to 4°C before use. Samples were ultracentrifuged at 36,000 rpm for 3h, 10 min, at 4°C, then samples were fractionated using a BioComp system, monitoring absorbance at 260 nm while collecting fractions of approximately 0.4 mL each.

### Oxidative phosphorylation

Real-time mitochondrial respiration was measured using the Seahorse XFe96 Extracellular Flux Analyzer (Agilent Technologies) and Seahorse XF Cell Mito Stress Test kit per manufacturer’s instructions. AML cells were treated with the indicated conditions prior to Seahorse assay and plated on Seahorse X96 plates coated with 20μL/well of Cell-TAK adhesive at 25μg/mL in 0.1M NaHCO3 at a concentration of 3x10^5^ cells/well suspended in XF base media supplemented with 1mM pyruvate, 2mM L-glutamine, 10mM glucose. OCR measurements were recorded with port injection of oligomycin A (1.5μM), FCCP (1μM), and rotenone/antimycin A (0.5μM) in order. Basal respiration and maximum respiration were calculated as follows. Basal respiration= (Last rate measurement before oligomycin A injection) - (Non-mitochondrial respiration rate), Maximal respiration= (Maximum rate measurement after FCCP injection)-(Non-mitochondrial respiration rate).

### EdU incorporation

10µM EdU was added to the cells 1 hour before harvest. EdU assays were performed using Click-iT™ EdU Alexa Fluor™ 488 Flow Cytometry Assay Kit (#C10420; Life Technologies) per manufacturer’s instructions. Cell cycle profiles were determined by flow cytometry (Millipore Guava easyCyte 8HT).

### Western blots

Western blot analyses were performed per manufacturer’s instructions using the following antibodies: ATM (A1106; Sigma Aldrich), p-ATM S1981 (ab91292; Abcam), cleaved caspase 3 (ab32042; Abcam), mTOR(#2972 and #2983; Cell Signaling), p-mTOR S2448 (#2971; Cell Signaling), c-MYC (#5605; Cell Signaling), p-H2AX S139 (#9718; Cell Signaling), p-4E-BP1 T37/47 (#2855; Cell Signaling), p-4E-BP1 S65 (#9456; Cell Signaling), p-S6 S240/244 (#2215; Cell Signaling), H3(#9715; Cell Signaling). p-AKT S473 (#4060; Cell Signaling), AKT (#9272; Cell Signaling). Each blot was quantified via densitometry by using Image Lab Software (Bio-Rad). Quantification data is shown in supplementary table.

### Therapeutic modeling in mice

NOD.Cg-Rag1^tm1Mom^II2rg^tm1Wj/^Tg(CMV-IL3,CSF2,KITLG)1Eav/J (NRG-S) mice were bred in-house. The patient samples for xenografting (from Dr. Daniel Pollyea, University of Colorado, Aurora, CO) were obtained from a 69-year-old female who was newly diagnosed with AML harboring FLT3-ITD and NPM1 mutations (AML #1, sample ID: HTB0097) and a 37-year-old female who developed relapse AML after chemotherapy (daunorubicin+cytarabine) with FLT3-ITD mutation (AML #2, sample ID: AML100510). Following expansion *in vivo,* as previously reported (32), the secondary leukemia was subsequently used for experiments. 24 hours prior to leukemic transplantation, 6-10 weeks old female NRG-S mice were conditioned with 25mg/kg busulfan delivered by i.p injection. Leukemia cells (1x10^6^) were injected i.v., and treatment started when peripheral blast counts were between 2% and 5% (mean 3.8%).

Leukemic burden was monitored by flow cytometry staining for human HLA-ABC-PE-Cy7 and CD45-FITC (eBioscience). Quizartinib was synthesized and prepared as previously described (32), and was dissolved in 30% Polyethylene glycol-400, 5% Tween 80 in ddH2O. 2.5mg/kg or 5mg/kg quizartinib was delivered once daily p.o. Everolimus was purchased from LC Laboratories, and was dissolved in the same solvent as quizartinib. 10mg/kg everolimus was delivered once daily p.o. After treatments, mice were sacrificed, followed by collection of leukemia cells harvested from the blood, spleen, and BM. Leukemic burden was measured by counting percentages of HLA-ABC+/CD45+ via flow cytometry. For RNA-seq analysis, human leukemia cells were isolated using EasySep^TM^ Mouse/Human Chimera Isolation Kit (#19849; Stemcell Technologies). All animal experiments were approved by and performed in accordance with guidelines of the Institutional Animal Care and Use Committee at University of Colorado (protocol No. 00170). Deidentified primary AML studies were obtained with donor consent from patients at the University of Colorado Anschutz Medical Campus (COMIRB protocol #12-0173).

### RPPA

MOLM-13 cells were treated with vehicle or 3nM quizartinib in either RPMI or hBMSC-CM for 16 hours, followed by washing with PBS. Cell pellets were frozen and sent to Reverse Phase Protein Array (RPPA) Core Facility at the University of Texas MD Anderson Cancer Center.

### RNA-seq

Following drug treatments and cell isolation from *in vitro* and *in vivo* studies as described above, RNA from each sample was isolated using RNeasy Plus Mini Kit (#73134; Qiagen). Poly-A-selected total RNA library construction was performed using Universal Plus mRNA-seq with NuQuant (#0520-A01;Tecan), and paired-end sequencing was performed on a NovaSeq6000 instrument (Illumina) in the Genomics Shared Resource (University of Colorado). Illumina adapters were trimmed and reads <50 base pairs were removed with BBDuk (57). Trimmed reads were aligned to the human Ensembl genome (hg38.p12, release 96) and gene counts were quantified using STAR v2.6.0a (58). Ensembl IDs were mapped to gene names and counts of genes with multiple IDs were aggregated. Lowly expressed genes were defined as <1 mean raw count or < 1 count per million across the dataset and were removed from the analysis. The limma R package (59) was used to calculate differential expression between the indicated groups. An interaction model was used with the input fractions for the polysome comparisons. Pathway analysis was performed using fold-change and the fgsea R package (60), and Hallmark or GO terms from the Molecular Signatures Database (61), which were downloaded using the msigdbr R package (62). Heatmaps were generated using the complexheatmap R package (63) following counts per million normalization and z-score transformation. Raw and processed data has been deposited to the Gene Expression Omnibus (GEO) – GEO identifier is pending.

### Public analysis of microarray data

Normalized expression was downloaded directly from GEO (GSE61019). AT case samples were compared to control using limma and pathway analysis was performed as described above.

### Statistical Analysis

Data are represented as box plots showing median values, with the boundaries of the rectangle representing the first and third quartiles, while whiskers extend from the minimum to maximum points in each plot. The number of replicates (n) is reported in the figure legends. Comparisons between 2 values were performed by Student t test (unpaired 2-tailed). Significance was defined at *P ≤ 0.05, **P ≤ 0.01, ***P ≤ 0.001, ****P ≤ 0.0001.

## Authors’ Contributions

**H.J. Park:** Conceptualization, data curation, data analysis, validation, investigation, visualization, methodology, writing–original draft, writing–review and editing. **M.A. Gregory:** Conceptualization, validation, investigation, methodology, writing-review and editing. **V. Zaberezhnyy:** Investigation, data curation**. A. Goodspeed:** Data analysis. **C.T. Jordan**: resources, conceptualization, writing-review **J.S. Kieft**: data curation, investigation, methodology. **J. DeGregori**: Supervision, conceptualization, resources, funding acquisition, validation, visualization, methodology, project administration, writing–review and editing.

## Acknowledgments

This work is supported by grants from NCI NRSA F30CA23197 (to H.J. Park), R35GM118070 (to J.S. Kieft), the V Foundation (T2016-012) and St. Baldrick’s Foundation (AWD-430131) (to J. DeGregori), and the Leukemia and Lymphoma Society (7020-19) (to C.T. Jordan and J. DeGregori). The authors thank Dr. Daniel Pollyea for providing human primary samples, Dr. Brett Stevens for critical reading of the manuscript, and the Bioinformatics and Biostatistics, Genomics and Functional Genomics Shared Resources supported by National Cancer Institute grant P30CA046934 to the University of Colorado Cancer Center. The authors thank the RPPA Core Facility at the University of Texas MD Anderson Cancer Center (NCI CA16672 and NIH R50CA22165).

---

**Figure S1.**
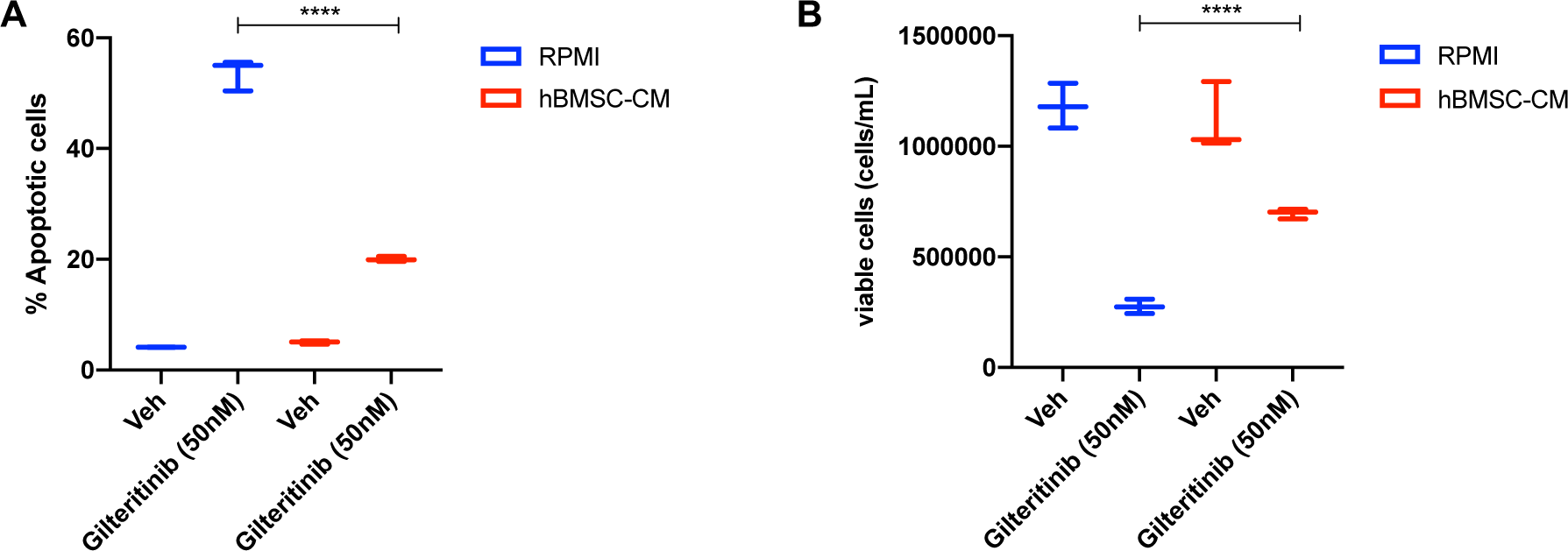
hBMSC-CM mediates protection against apoptosis in FLT3-ITD AML cells following FLT3 inhibition by gilteritinib. **A** and **B**, MOLM-13 cells were treated with either DMSO (veh) or 50nM gilteritinib for 48 hours in RPMI or hBMSC-CM, followed by measurement of (**A**) apoptosis and (**B**) cell viability (n=3).

**Figure S2.**
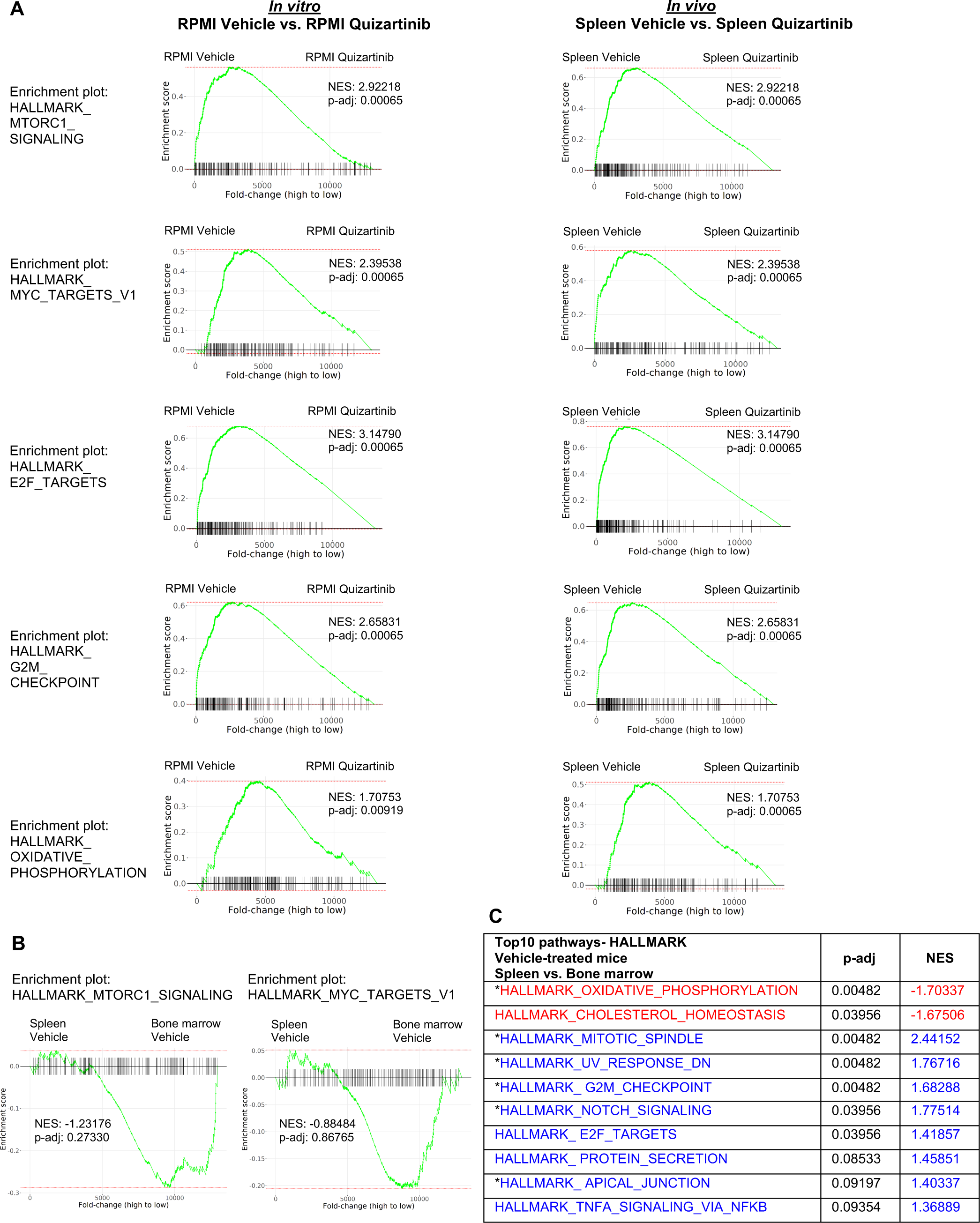
Similar pathways are altered in FLT3-ITD AML cells following FLT3 inhibition *in vitro* and *in vivo* in the absence of bone marrow microenvironment. **A,** GSEA of Hallmark gene sets was performed to determine the effect of FLT3 inhibition in the absence of BM microenvironment. *In vitro*: RNA-seq data from vehicle-treated cells in RPMI was compared with those from quizartinib-treated cells in RPMI. *In vivo*: RNA-seq data from samples from the spleen of vehicle-treated mice was compared with those from the spleen of quizartinib-treated mice. Enrichment plots of 5 most significant pathways from each type of study are shown. **B**, Enrichment plots showing mTORC1 signaling or MYC targets in AML cells isolated from the spleen compared to those from the BM of vehicle-treated mice. **C**, Top 10 enriched pathways (by p-adj) from GSEA of Hallmark gene sets applied on RNA-seq data from human AML cells. Samples from the spleen of vehicle-treated mice were compared with the samples from the BM of the same paired mouse. Pathways with significant alteration (p-adj < 0.1) are represented with colors (Red: negative NES, Blue: positive NES). Asterisks represent matched pathways with same enrichment patterns observed from GSEA of Hallmark gene sets comparing the spleen from quizartinib-treated mice with BM of the same paired mouse (Fig.2G).

**Figure S3.**
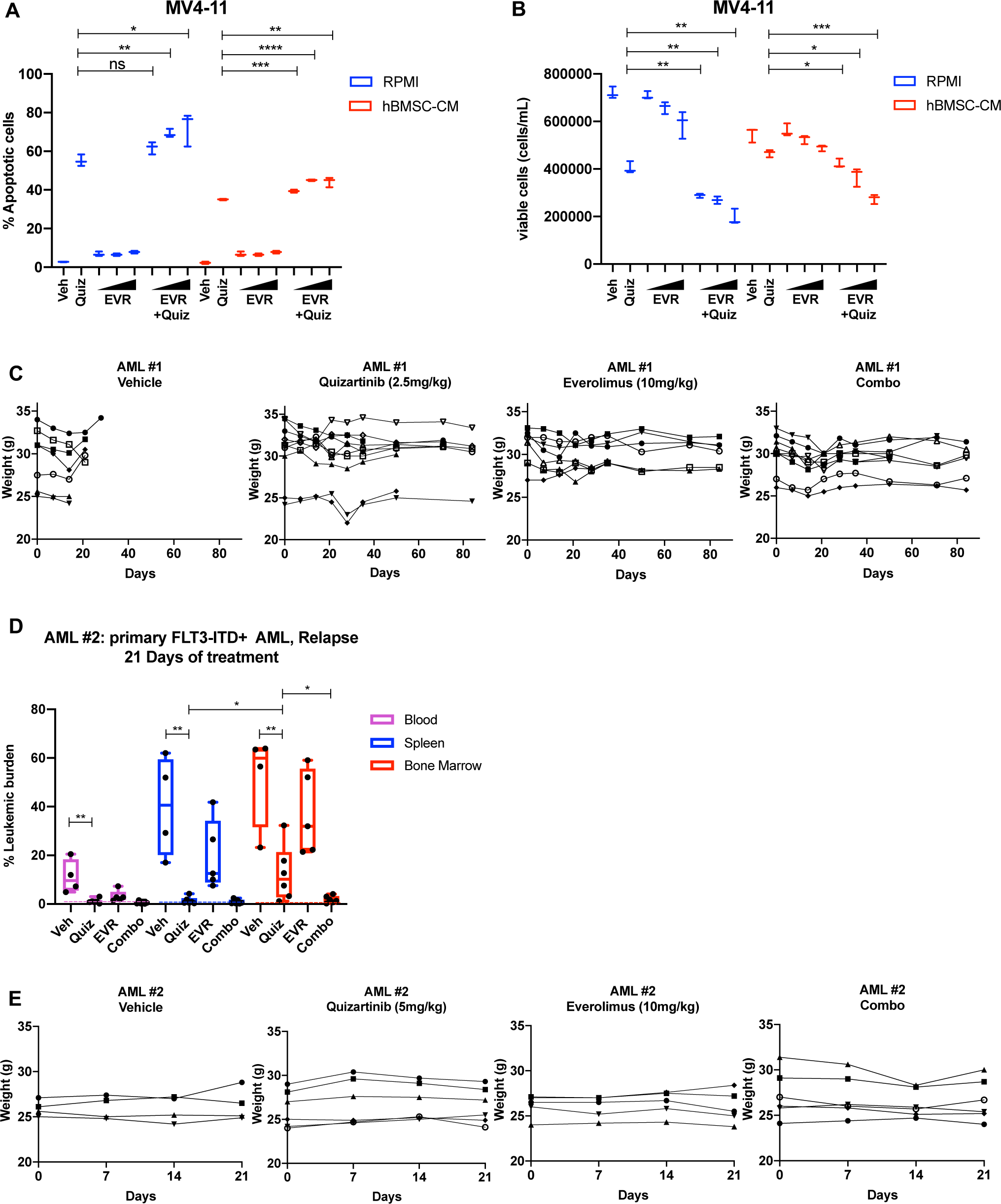
Combinatorial treatment of quizartinib with everolimus demonstrates significantly better cell killing effects *in vitro* and does not show major signs of toxicity *in vivo*. A and B, MV4-11 cells were treated with 2nM quizartinib (Quiz) and everolimus (EVR) alone or together as indicated (increasing concentrations of everolimus of 5, 10, and 50nM indicated by triangles) for 48 hours in RPMI or hBMSC-CM (n=3), followed by measurement apoptosis (A) and cell viability (B) as previously described. C, Mice weight from each group of long-term treatment of primary AML #1 was measured weekly. Throughout the duration of long-term treatment, vehicle-treated mice had a median survival of 27 days. A few mice across other groups were found to be dead without signs of evident leukemia or weight loss. D and E, NRG-S mice were transplanted with human primary FLT3-ITD^+^ AML (AML #2: a 37-year-old female patient who developed relapse after chemotherapy). When leukemic burden reached 2- 5% (∼4 months after transplantation, determined by measuring CD45^+^ HLA-ABC^+^ cells in the blood), mice were treated with 1) vehicle (Veh), 2) 5mg/kg quizartinib (Quiz), 3) 10mg/kg everolimus (EVR), 4) combo for 21 days, daily p.o. (Veh: n=4, Quiz: n=6, EVR: n=5, Combo: n=6). After treatment, leukemic burden was measured by quantifying HLA-ABC^+^ CD45^+^ cells via flow cytometry in spleen, BM, and blood. Dotted line represents negative controls from non-transplanted mice. (D) Plot showing leukemic burden of each treatment group; dotted line represents negative controls from non-transplanted mice. (E) Mice weight from each group of treatment of primary AML #2 was measured weekly.

**Figure S4.**
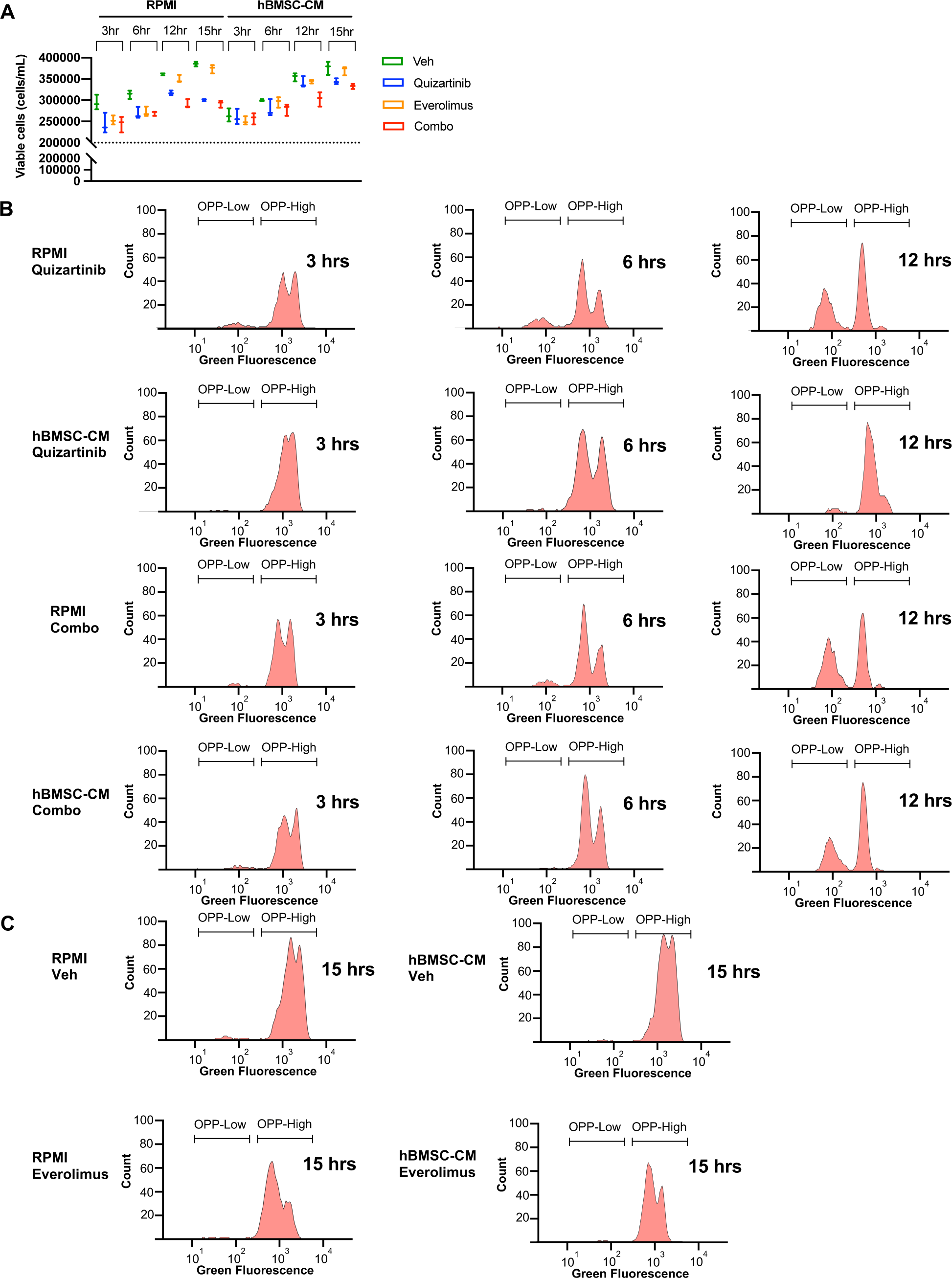
FLT3-ITD AML cells demonstrate shift of OPP-labeled peaks over the course of drug treatments without showing signs of cell death. A, Cell viability was measured at each time condition prior to OPP assay. Cell density at the start of treatment (2x10^5^ cells/mL) is represented as dotted line. B and C, Representative flow cytometry histograms of the OPP assay from (B) cells treated with quizartinib or quizartinib/everolimus combo for shorter timepoints (3, 6, and 12 hours) either in RPMI or hBMSC-CM, and (C) cells treated with vehicle or everolimus only for 15 hours either in RPMI or hBMSC-CM.

**Figure S5.**
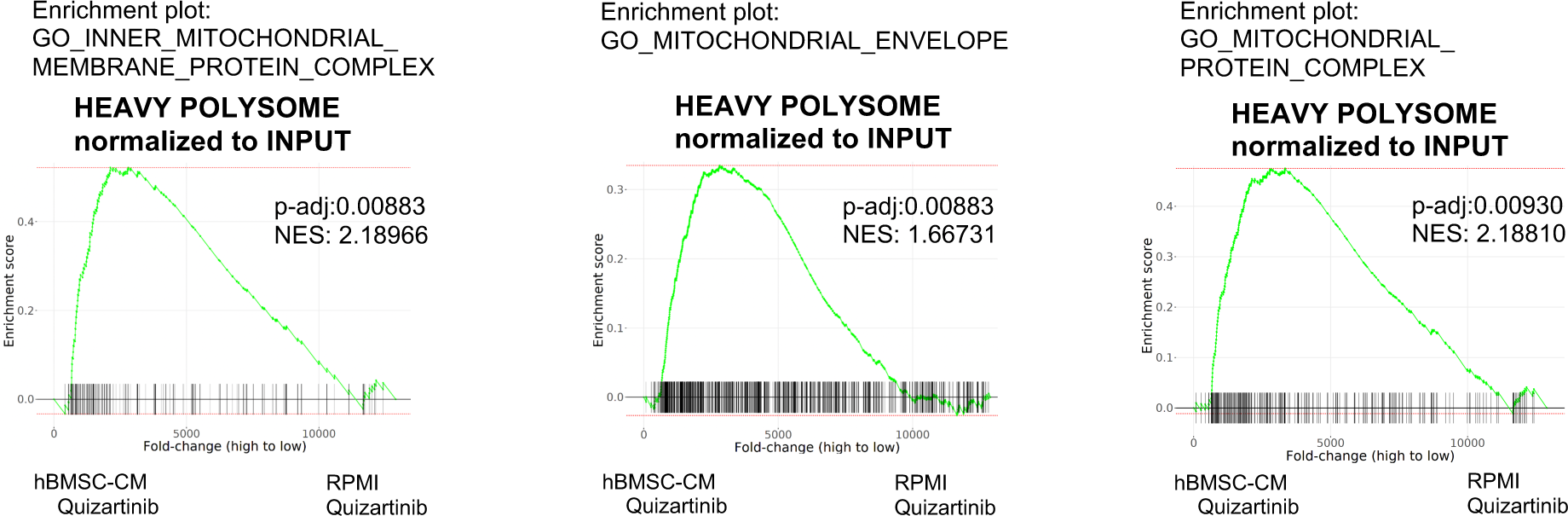
Transcripts associated with mitochondrial integrity are enriched in polysomes of FLT3-ITD AML cells in hBMSC-CM upon FLT3 inhibition. GSEA of Gene Ontology gene sets applied on RNA-seq data from human AML cells from heavy polysome-associated mRNAs normalized to input. Pathways associated with mitochondrial integrity are shown.

**Figure S6.**
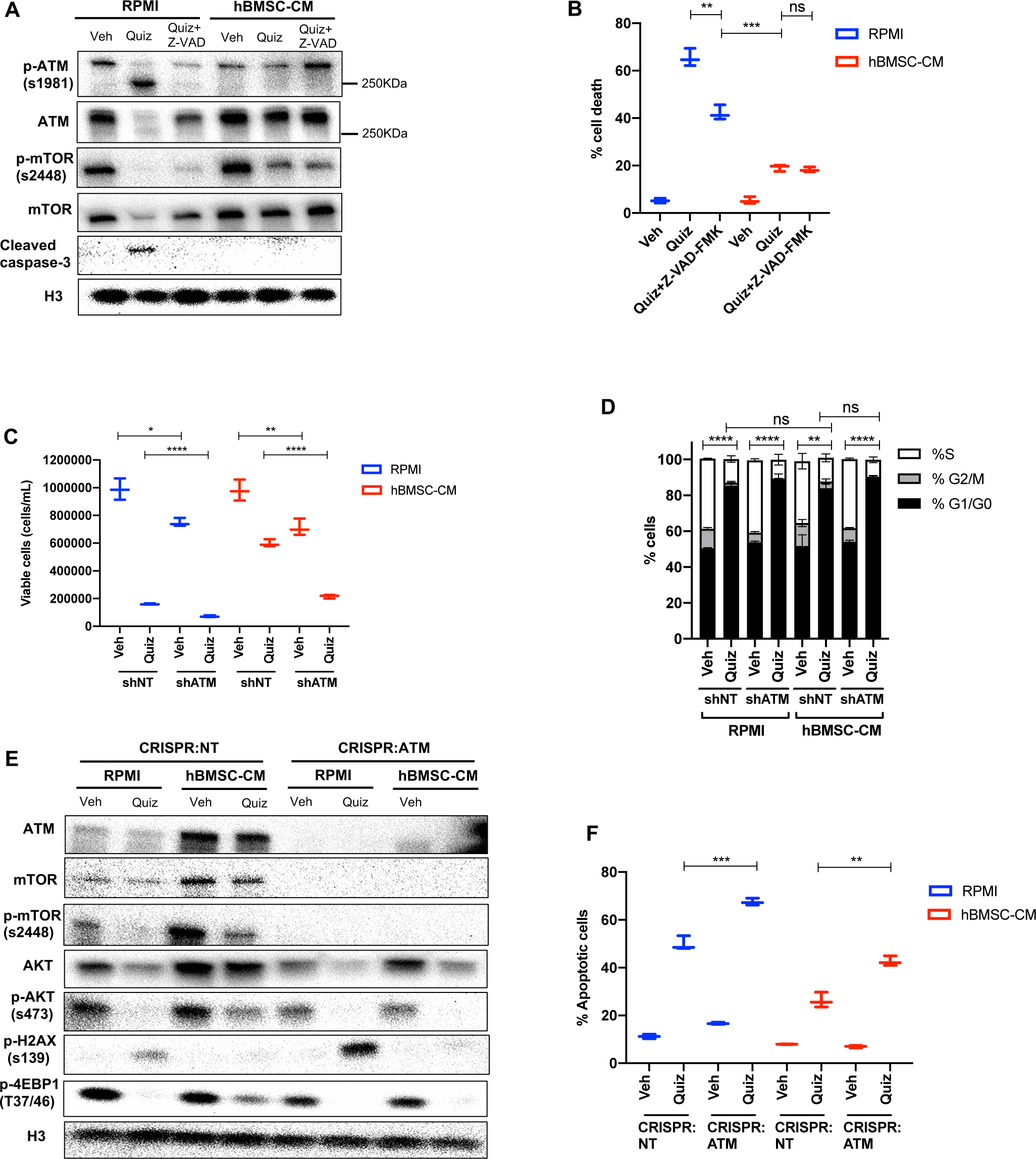
ATM controls mTOR expression and AKT phosphorylation without affecting the cell cycle. **A** and **B**, MOLM-13 cells were treated with vehicle, 3nM quizartinib (Quiz), and 3nM quizartinib+60uM Z-VAD-FMK (Z-VAD) in RPMI or hBMSC-CM for (**A**)15 hours and harvested to determine levels of the indicated proteins by Western blotting and (**B**) 48 hours, followed by measurement of cell death by PI staining (n=3). **C**, MOLM-13 cells were transduced with lentiviruses expressing shRNAs (shNT: non-targeting shRNA or shATM: ATM-targeting shRNA). Cells were treated with either vehicle or 3nM quizartinib (Quiz) for 48 hours in RPMI or hBMSC-CM, followed by measurement of cell viability (n=3). **D**, MOLM-13 cells expressing shNT or shATM were treated with the indicated conditions for 15 hours and labeled with 5-Ethynyl-2′-deoxyuridine (EdU). Cells were stained with PI to measure DNA content, followed by cell cycle analysis using flow cytometry (S-phase: EdU+, G1/G0-phase: EdU- /low PI, G2/M-phase: EdU- /high PI) (n=3). Statistical analysis was performed for S phase. **E** and **F**, MOLM-13 cells were transduced with lentiviruses expressing CRISPR/Cas9 with sgRNA (CRISPR:NT; non-targeting and CRISPR:ATM; ATM-targeting sgRNA), followed by plating on methylcellulose for single cell-derived clone isolation. After expansion of single cells, the cells were treated with either vehicle or 3nM quizartinib (Quiz) for (**E**)15 hours and harvested to determine levels of the indicated proteins or (**F**) 48 hours, followed by measurement of apoptosis (n=3).

**Figure S7.**
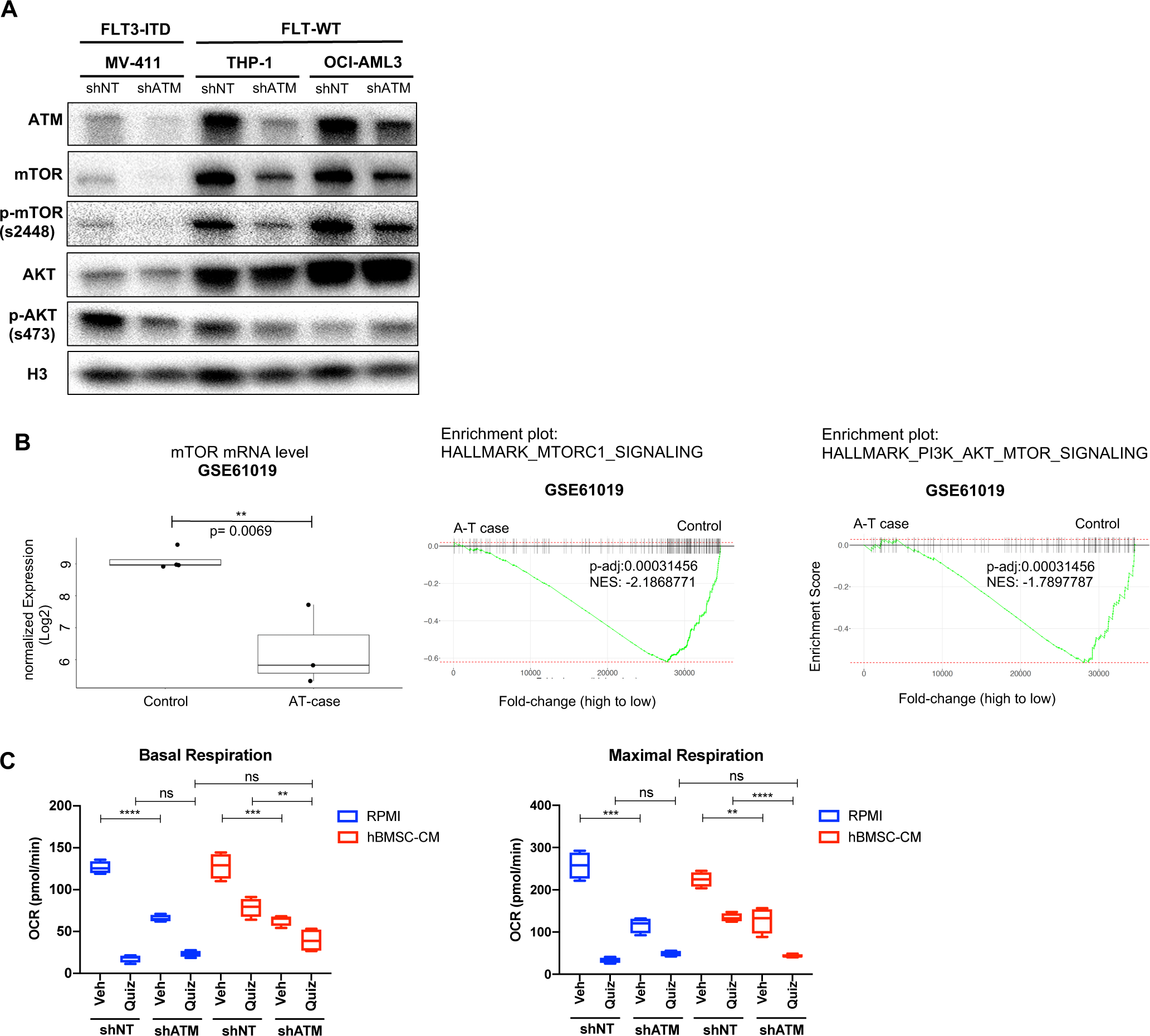
ATM-dependent mTOR expression and AKT phosphorylation are not unique to FLT3-dependent AML or leukemic cells in general. **A**, MV4-11, THP-1, and OCI-AML3 cells expressing shNT or shATM were harvested to determine levels of the indicated proteins by Western blot analysis. **B**, Analysis of RNA-seq data obtained from GEO database (GSE61019). Level of MTOR mRNAs from human brain tissues of A-T cases were compared with those from control samples (left) and GSEA enrichment plots of mTORC1 signaling and PI3K-AKT-mTOR signaling are shown. **C**, MOLM-13 cells expressing shNT or shATM were treated with the indicated conditions for 15 hours, followed by measurement of OCR by Seahorse XF Cell Mito Stress Test (n=4). Basal respiration and maximal respiration are shown.

**SUPPLEMENTARY TABLE.**

| **Quantification of Western blot images by densitometry**<br>**All values were normalized to RPMI Veh**<br>**Average values of biological replicates are shown (n=2-3)** |  |  |  |  |  |  |  |  |  |  |
| --- | --- | --- | --- | --- | --- | --- | --- | --- | --- | --- |
| **Fig. 1C** | **RPMI Veh** | **RPMI Quiz** | **hBMS C-CM Veh** | **hBMS C-CM Quiz** |  |  |  |  |  |  |
| **pFLT3 (Y593)/Tubulin** | 1 | 0.094 | 0.812 | 0.087 |  |  |  |  |  |  |
| **FLT3/Tubulin** | 1 | 1.663 | 0.790 | 2.632 |  |  |  |  |  |  |
| **Fig. 2D** | **RPMI Veh** | **RPMI Quiz 3hr** | **RPMI Quiz 6hr** | **RPMI Quiz 10hr** | **RPMI Quiz 15hr** | **hBMSC -CM Veh** | **hBMSC -CM Quiz 3hr** | **hBMSC -CM Quiz 6hr** | **hBMSC -CM Quiz 10hr** | **hBMSC -CM Quiz 15hr** |
| **p-mTOR (s2448)/H3** | 1 | 0.852 | 0.312 | 0.331 | 0.211 | 1.832 | 1.616 | 1.412 | 1.212 | 1.352 |
| **p-S6 (s240/s244)/H3** | 1 | 0.312 | 0.218 | 0.359 | 0.419 | 1.235 | 0.415 | 0.661 | 0.552 | 0.691 |
| **c-MYC/H3** | 1 | 0.274 | 0.316 | 0.221 | 0.322 | 1.632 | 0.412 | 0.492 | 0.789 | 0.762 |
| **Fig. 4B** | **RPMI Veh** | **RPMI Quiz** | **hBMS C-CM Veh** | **hBMS C-CM Quiz** | **hBM SC-CM EVR** | **hBMSC -CM Combo** |  |  |  |  |
| **p-mTOR (s2448)/H3** | 1 | 0.197 | 1.122 | 0.530 | 0.444 | 0.237 |  |  |  |  |
| **mTOR/H3** | 1 | 0.768 | 1.340 | 0.986 | 0.862 | 0.799 |  |  |  |  |
| **p-S6 (s240/s244)/H3** | 1 | 0.274 | 1.370 | 0.506 | 0.136 | 0.075 |  |  |  |  |
| **p-4E-BP1 (s65)/H3** | 1 | 0.076 | 1.412 | 0.481 | 0.496 | 0.087 |  |  |  |  |
| **p-4E-BP1 (T37/46)/H3** | 1 | 0.182 | 1.884 | 0.301 | 0.641 | 0.246 |  |  |  |  |

| **Fig. 7A** | **RPMI Veh** | **RPMI Quiz** | **hBMS C-CM Veh** | **hBMS C-CM Quiz** | **hBM SC-CM EVR** | **hBMSC -CM Combo** |  |  |
| --- | --- | --- | --- | --- | --- | --- | --- | --- |
| **ATM/H3** | 1 | 0.562 | 1.206 | 0.851 | 1.160 | 0.775 |  |  |
| **p-ATM (s1981) upper band 350KDa /H3** | 1 | 0.484 | 1.510 | 1.312 | 2.045 | 1.078 |  |  |
| **p-ATM (s1981) lower band 250KDa /H3** | 1 | 10.33 | 0.598 | 2.412 | 1.234 | 4.7240 |  |  |
| **Fig. 7C** | **shNT RPMI Veh** | **shNT RPMI Quiz** | **shNT hBMS C-CM Veh** | **shNT hBMS C-CM Quiz** | **shAT M RPMI Veh** | **shATM RPMI Quiz** | **shATM hBMSC -CM Veh** | **shATM hBMSC -CM Quiz** |
| **ATM/H3** | 1 | 0.969 | 2.053 | 1.182 | 0.043 | 0.039 | 0.049 | 0.093 |
| **mTOR/H3** | 1 | 1.159 | 2.436 | 2.111 | 0.108 | 0.103 | 0.164 | 0.162 |
| **p-mTOR (s2448)/H3** | 1 | 0.021 | 2.145 | 0.792 | 0.059 | 0.036 | 0.041 | 0.031 |
| **AKT/H3** | 1 | 0.967 | 1.259 | 1.162 | 0.541 | 0.284 | 0.617 | 0.542 |
| **p-AKT (s473)/H3** | 1 | 0.319 | 2.596 | 0.639 | 2.047 | 0.217 | 1.715 | 0.271 |
| **pH2AX(s139)/H3** | 1 | 5.652 | 0.674 | 1.210 | 1.240 | 7.991 | 0.596 | 0.555 |
| **p-4E-BP1 (T37/46)/H3** | 1 | 0.342 | 2.115 | 1.312 | 1.815 | 0.279 | 1.412 | 0.415 |
| **p-4E-BP1 (s65)/H3** | 1 | 0.141 | 2.234 | 0.810 | 2.155 | 0.212 | 1.920 | 0.254 |
| **p-S6(s240/244)/H3** | 1 | 0.249 | 2.412 | 0.781 | 1.231 | 0.211 | 1.123 | 0.192 |
| **Fig. S6A** | **RPMI Veh** | **RPMI Quiz** | **RPMI Quiz+ Z-VAD** | **hBMS C-CM Veh** | **hBM SC-CM Quiz** | **hBMSC -CM Quiz+ Z-VAD** |  |  |
| **p-ATM (s1981) upper band 350KDa /H3** | 1 | 0.216 | 0.516 | 0.782 | 0.611 | 1.634 |  |  |
| **p-ATM (s1981) lower band 250KDa /H3** | 1 | 9.163 | 1.155 | 0.823 | 0.724 | 0.823 |  |  |
| **ATM/H3** | 1 | 0.277 | 0.842 | 1.072 | 0.660 | 0.862 |  |  |
| **p-mTOR (s2448)/ H3** | 1 | 0.085 | 0.189 | 1.190 | 0.556 | 0.512 |  |  |
| **mTOR/ H3** | 1 | 0.345 | 0.738 | 1.075 | 0.936 | 0.980 |  |  |
| **Cleaved caspase-3 /H3** | 1 | 9.155 | 0.851 | 0.912 | 1.051 | 0.725 |  |  |
| **Fig. S6E** | **crispr :NT RPMI Veh** | **crispr :NT RPMI Quiz** | **crispr :NT hBMS C-CM Veh** | **crispr :NT hBMS C-CM Quiz** | **crispr :ATM RPMI Veh** | **crispr :ATM RPMI Quiz** | **crispr :ATM hBMS C-CM Veh** | **crispr :ATM hBMS C-CM Quiz** |
| **ATM/H3** | 1 | 0.721 | 3.153 | 2.414 | 0.023 | 0.049 | 0.071 | 0.214 |
| **mTOR/ H3** | 1 | 0.862 | 2.152 | 2.013 | 0.212 | 0.313 | 0.251 | 0.215 |
| **p-mTOR (s2448)/ H3** | 1 | 0.041 | 2.891 | 0.690 | 0.049 | 0.026 | 0.053 | 0.040 |
| **AKT/H3** | 1 | 0.767 | 1.821 | 1.362 | 0.841 | 0.271 | 1.152 | 0.501 |
| **p-AKT (s473)/ H3** | 1 | 0.135 | 1.324 | 0.639 | 0.739 | 0.192 | 0.715 | 0.291 |
| **pH2AX( s139) /H3** | 1 | 4.824 | 1.523 | 1.120 | 1.240 | 9.133 | 0.896 | 1.555 |
| **p-4E-BP1 (T37/46) /H3** | 1 | 0.209 | 0.819 | 0.352 | 0.592 | 0.199 | 0.668 | 0.212 |
| **Fig. S7A** | **MV4-11 shNT** | **MV4-11 shATM** | **THP-1 shNT** | **THP-1 shATM** | **OCI-AML3 shNT** | **OCI-AML3 shATM** |  |  |
| **ATM/H3** | 1 | 0.397 | 1 | 0.436 | 1 | 0.552 |  |  |
| **mTOR/**<br>**H3** | 1 | 0.297 | 1 | 0.498 | 1 | 0.714 |  |  |
| **p-**<br>**mTOR**<br>**(s2448)/**<br>**H3** | 1 | 0.519 | 1 | 0.499 | 1 | 0.717 |  |  |
| **AKT/H3** | 1 | 1.093 | 1 | 0.936 | 1 | 0.942 |  |  |
| **p-AKT**<br>**(s473)/**<br>**H3** | 1 | 0.527 | 1 | 0.742 | 1 | 1.594 |  |  |

